# Monoclonal antibody chP3R99 reduces subendothelial retention of atherogenic lipoproteins in Insulin-Resistant rats: *Acute treatment versus long-term protection as an idiotypic vaccine for atherosclerosis*

**DOI:** 10.1101/2023.08.30.555546

**Authors:** Yosdel Soto, Arletty Hernández, Roger Sarduy, Victor Brito, Sylvie Marleau, Donna F. Vine, Ana M. Vázquez, Spencer D. Proctor

**Author notes:** **Joint Corresponding Authors:** Dr. Spencer D. Proctor, 4-002J Li Ka Shing Centre, University of Alberta, Edmonton, Alberta, Canada. T6G2P5. Email address. Dr. Yosdel Soto. Centre for Molecular Immunology, 216 St and 15th Ave, Atabey, Playa PO Box 16040, Havana 11600, Havana, Cuba.

## Abstract

**BACKGROUND:** Atherosclerosis is triggered by the retention of apolipoprotein B-containing lipoproteins by proteoglycans. In addition to LDL, remnant lipoproteins have emerged as pivotal contributors to this pathology, particularly in the context of insulin resistance and diabetes. We have previously reported anti-atherogenic properties of a monoclonal antibody (chP3R99) that recognizes sulfated glycosaminoglycans on arterial proteoglycans.

**METHODS AND RESULTS:** Solid-phase assays demonstrated that chP3R99 effectively blocked over 50% lipoprotein binding to chondroitin sulfate and vascular extracellular matrix *in vitro*. The pre-perfusion of chP3R99 (competitive effect) resulted in specific antibody-arterial accumulation and reduced fluorescent lipoprotein retention by ∼60% in insulin resistant JCR:LA-*cp* rats. This competitive reduction was dose-dependent (25 µg/mL–250 µg/mL), effectively decreasing deposition of cholesterol associated with lipoproteins. In a five-week vaccination study in insulin resistant rats with (200 µg SC, once a week), chP3R99 reduced arterial lipoprotein retention, and was associated with the production of anti-chondroitin sulfate antibodies (Ab3) able to accumulate in the arteries (dot-blot). Neither the intravenous inoculation of chP3R99 (4.5 mg/kg), nor the immunization with this antibody displayed adverse effects on lipid or glucose metabolism, insulin resistance, liver function, blood cell indices, or inflammation pathways in JCR:LA*-cp* rats.

**CONCLUSIONS:** Both acute (passive) and long-term administration (idiotypic cascade) of chP3R99 antibody reduced LDL and remnant lipoprotein interaction with proteoglycans in an insulin-resistant setting. These findings support the innovative approach of targeting pro-atherogenic lipoprotein retention by chP3R99 as a passive therapy or as an idiotypic vaccine for atherosclerosis.

**CLINICAL PERSPECTIVE:** *What Is New?:*

- Innovative anti-atherosclerotic chP3R99 mAb interferes with proteoglycan binding of both LDL and remnant lipoproteins *in vitro* and *in vivo*.
- *In vivo* kinetic studies that immunize with chP3R99 reveal a temporal induction of an anti-idiotypic antibody cascade in a model of insulin resistance (analogous to a vaccine).
- We discovered that the idiotypic chP3R99 monoclonal antibody (Ab1) was able to induce protective anti-anti-idiotypic (Ab3) antibodies present in both sera as well as the aorta (target organ) *in vivo*.

*What Are The Clinical Implications?:*

- The chP3R99 mAb has efficacy for reducing the arterial retention of both LDL and remnant-derived lipoproteins and may be relevant of those with Type-2 Diabetes and/or residual CVD risk.
- We show efficacy of chP3R99 mAb under pro-inflammatory conditions and that it does not exacerbate other metabolic aberrations during insulin resistance.
- Data support the targeting pro-atherogenic lipoprotein retention with chP3R99 as a passive therapy or as an idiotypic vaccine for atherosclerosis, complementary to lipid lowering approaches.

## INTRODUCTION

Atherosclerotic vascular disease (ASVD) afflicts millions of individuals each year, particularly those with insulin-resistance or diabetes (1,2). Despite the rapid discovery of new cholesterol and lipid lowering agents, the continued rise in the incidence of ASVD suggests that we are yet to completely understand the etiology of atherosclerosis. The ‘response-to-retention’ hypothesis for atherosclerosis suggests that lipid and cholesterol accumulate in arterial vessels derived from fasting and non-fasting lipoproteins (3–4). Atherogenic cholesterol-dense lipoproteins (for example low density lipoprotein, LDL) are thought to permeate both intact and/or damaged arterial endothelium, become entrapped within the sub-endothelial space and accumulate, resulting in inflammation and atheroma (3–4). Although the literature documents a significant epidemiological and/or genetic (i.e. GWAS) association between raised circulating fasting LDL cholesterol (LDL-C) and ASVD risk, a large proportion of subjects diagnosed with ASVD are either normolipidemic (normal levels of LDL-C), or have substantial residual risk (5–7). These data suggest that clinically, atheromata-associated cholesterol is derived from alternate sources, including non-fasting remnant lipid fractions (8).

In support of this concept, a series of publications authored by the Copenhagen Heart Study group (as well as others) have provided evidence for a major shift in the paradigm of atherogenesis (and potentially its therapy), suggesting that remnant (non-fasting) cholesterol is a major causative factor for ASVD (9–13). These clinical outcomes have caused us to refine our focus to how we might therapeutically target remnant cholesterol. Traditional therapeutic strategies have focused on the modulation of risk factors for ASVD (i.e. lipid lowering) rather than halting lipoprotein retention in the artery wall *per se.* Importantly, there have been key contributions to this field (by us and others) (14–19), that provide proof of principal that targeting the interaction of both LDL and remnant lipoproteins specifically within the arterial wall is a unique and viable therapeutic strategy to reduce atherosclerosis.

Histological examination of arterial proteoglycan distribution in human tissue with ASVD has demonstrated increases in chondroitin sulfate (CS) proteoglycans including biglycan, decorin and versican, compared to vessels without disease (20,21). The literature also documents that the capacity for proteoglycan-associated glycosaminoglycans (GAGs) to bind LDL (as well as remnants), is related to the electrostatic properties and length of their side chains (22–27).

To complicate matters, we also now appreciate that the condition of insulin resistance is a critical precursor to Type-2 diabetes with the sequelae to ASVD (27–30). We know that chronic insulin resistance results in changes in arterial architecture, vascular perturbations, and dyslipidemia. Vascular dysfunction is associated with insulin resistance can re-model (stimulate) extracellular arterial CS-proteoglycans and intimal thickening (27–30). Excessive proliferation of vascular smooth muscle cells (VSMC) can stimulate the secretion of arterial CS-proteoglycans, which in turn increases the capacity for lipoprotein binding (27–32). Several lines of evidence suggest that this produces a unique sulfation pattern on arterial GAGs during ASVD, propagated by the tumor growth factor-β (33–36).

The interaction of atherogenic lipoproteins with arterial proteoglycans offers an explicit target for atherogenesis, but the technological capacity for this approach has not been viable until now. For this study we proposed to utilize an innovative monoclonal antibody (mAb) chP3R99 (made possible by a collaboration with the Centro de Inmunología Molecular, Cuba). Investigators at Centro de Inmunología Molecular have pioneered the development of a unique mAb that recognizes sulfated GAGs found within CS-proteoglycans (37–40) (Figure 1).

**Figure 1.**
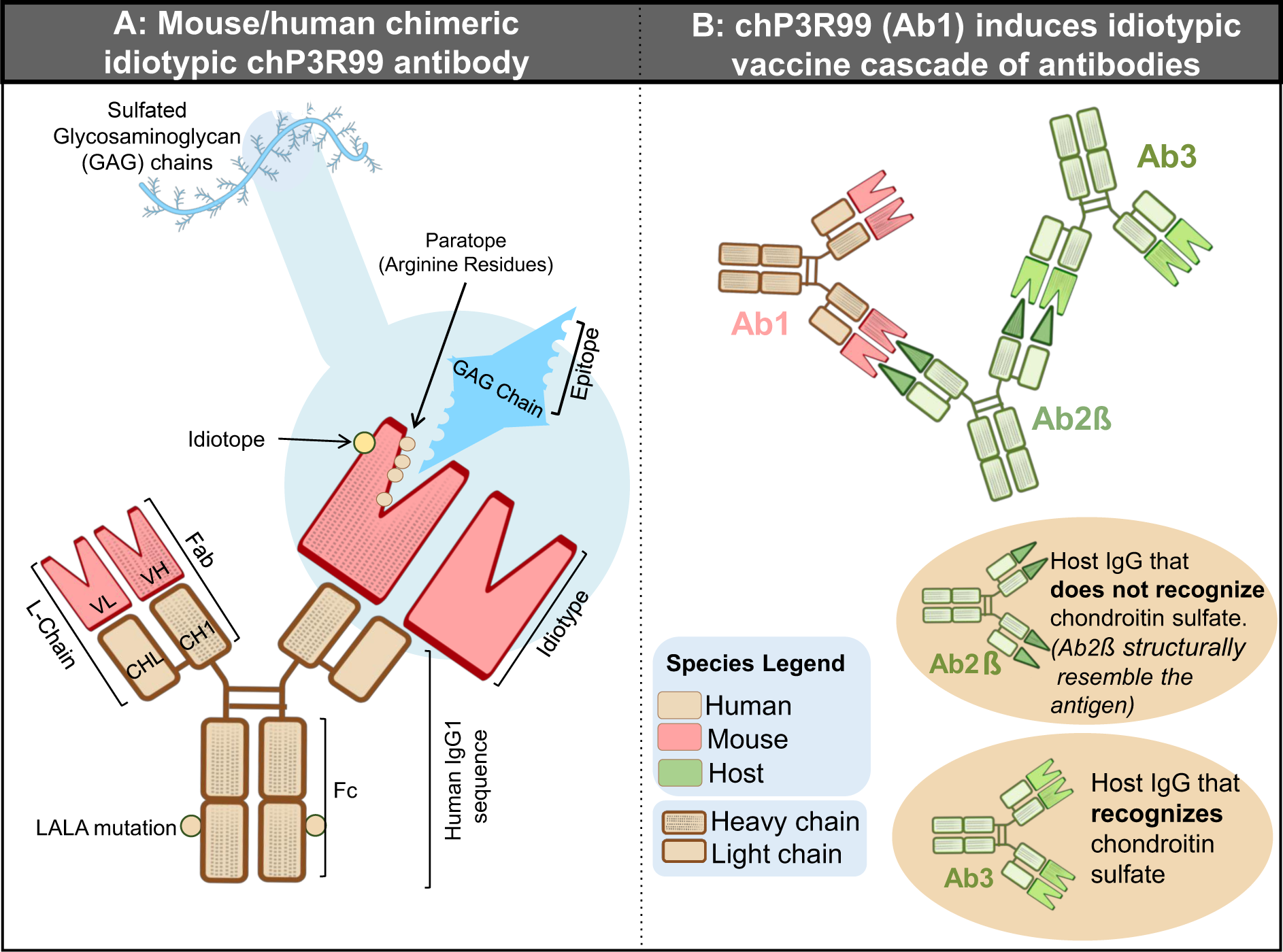
Summary of the main features of chP3R99 mAb. (A) Chimeric mAb composed by mouse variable regions (idiotype) and human IgG1constant regions mutated to abrogate the binding to Fcγ receptors and the complement system activation (LALA mutation). The antigen recognition site of chP3R99 is enriched in arginine residues involved in the ‘Acute’ (competitive) interaction with the lipoprotein binding site on glycosaminoglycan side chains of proteoglycans. The variable regions of chP3R99 contains immunogenic epitopes (idiotopes) including the antigen-combination site (paratopes). (B) Schematic of the putative idiotypic vaccine cascade induced by the host immunized with chP3R99 mAb (Ab1) to produce anti-idiotypic antibodies (Ab2) and anti-anti-idiotypic antibodies (Ab3).

Centro de Inmunología Molecular originally developed the murine P3 mAb devised to recognize N-glycosilated gangliosides and sulfatides (41). The P3 mAb was characterized with the unusual property of generating a strong anti-idiotypic response when administered in syngeneic animal models (even in the absence of adjuvants or carrier proteins) (42–44). Subsequently, a chP3 mutant was engineered with an additional arginine in position 99 at the Complementarity-determining Region 3 (H-CDR3) from the heavy chains (Kabat numbering; chP3R99), that displayed an even higher reactivity with antigens, especially sulfatides (45,46). Finally, the Fcγ receptor and complement-binding region of the chP3R99 mAb was modified to include the ‘LALA’ mutation. Interestingly, the LALA sequence impaired undesired induction of pro-inflammatory responses (from the Th1 infection pathway), while still maximising the antibody production response (from the Th2 B-cell pathway which is anti-inflammatory and associated with neutralizing Ab production) (Figure 1A) (37).

The chP3R99 mAb is known to interfere directly with the interaction of LDL to CS *in vitro* and markedly reduces the retention of LDL *in vivo* (37–40). Importantly, the chP3R99 has been shown to accumulate in lesion areas *in vivo* (47). These advancements suggest a high degree of specificity for CS within arterial lesions and reduced affinity for other enriched sites; suggesting specificity for a unique CS sulfation expressed on vascular GAGs (37–40). For this study we proposed that; (i) the chP3R99 mAb would interfere (acutely) with both LDL and remnant lipoproteins both *in vitro* and *in vivo*; and that (ii) immunizations with chP3R99 in rodent model of insulin resistance would similarly reduce overall retention of lipoproteins but be associated with the induction of 2^nd^ and 3^rd^ Ab responses consistent with an idiotypic vaccine cascade effect (48).

## METHODS

### Data availability

The data associated with the findings of this study are available from the corresponding author upon reasonable request. Supplementary method descriptions for Data S1-4 and Tables S1-6 can be found in the Supplementary Materials.

### Antibodies

ChP3R99 (Figure 1) and hR3 (anti-human epidermal growth factor receptor) mAbs were provided by the Center of Molecular Immunology (Havana, Cuba). As per previous published studies, the mAbs were purified by protein-A affinity chromatography (Pharmacia, Sweden) from transfectome culture supernatants and analyzed by SDS-PAGE under reducing conditions (37, 38). The specificity of the purified mAbs was confirmed by enzyme-linked immunosorbent assay (ELISA) and protein concentration was estimated by optical density at 280 nm. In all set of experiments, an Ab variant with impaired capacity to bind Fc gamma receptor (FcγR) and the complement system was used to avoid undesired Fc-mediated pro-inflammatory effects (chP3R99-LALA, referred to as chP3R99 subsequently).

### Animal Model

Experiments in animals were conducted in JCR:LA-cp (cp/cp) male rats (400-600 g), a spontaneously recessive strain with an absence of the leptin receptor leading to hyperphagia-associated obesity, along with severe insulin resistance. As a metabolically normal control, age-matched lean (+/cp) rats (350-400 g) from the same strain fed a chow diet throughout the studies were used. The rats were raised in our well-established colony at the University of Alberta (Edmonton, Canada) under standard conditions (25°C, 60±10% humidity), 12 h day/night cycles with water and food *ad libitum* (16). All experiments were conducted as per the Canadian Council for Animal Care and approved under the University of Alberta protocol numbers AUP000952 and AUP0004174. The rationale for the sample size for rat studies was based on previously published work with this strain and the ethically minimum number of animals required to detect a meaningful biologically relevant difference in treatment groups.

### Arterial Perfusion Experiments and General Study Design(s)

Acute perfusion experiments with the mAbs were conducted at 12 weeks of age in insulin resistant rats fed a chow diet as previously described (14–16). For subsequent experiments, 10-week-old insulin resistant rats were subject to a lipid-balanced diet supplemented with 1% w/w cholesterol for six weeks (49). Physiological studies to address potential toxicity of chP3R99 administration were conducted both in lean control and insulin resistant rats. In the acute experiments, the rats received a bolus IV injection of the mAbs at 4.5 mg/kg in PBS through the jugular vein at 15 weeks of age. Conversely, to assess the impact of the vaccination with this mAb, the rats were given six SC injections of P3R99 mAb (200 µg, once a week), hR3 isotype control or saline (n=6) along with the high fat diet feeding. In both settings, blood samples were drawn at different time points for further analysis in sera and plasma. Also, a complete blood count was carried out before starting the treatment with the mAbs and at the endpoint by mean of a Coulter STKS instrument (Coulter Electronics). The animals were fasted overnight, anesthetized under 3.5% isofluorane-oxygen for surgical procedures and sacrificed by exsanguination. Tissue samples were collected appropriately for histopathological, immunofluorescence, immunoblotting, and biochemical evaluations.

### Lipoprotein Preparations

Intestinal remnant lipoproteins were isolated and labeled with fluorophores following a methodology as previously described (14–16). Chylomicrons were labeled with Cy5 according to manufactures instructions (GE Healthcare UK limited, Buckinghamshire, UK) and injected into hepatectomized rabbits (n=3) to obtain remnants endogenously (14–16). The later were isolated from rabbit sera by ultracentrifugation, whereas human LDL were obtained commercially and labeled with Cy3 (Millipore Sigma Canada Ltd, ON, Canada). Previous studies have demonstrated that ApoB48-Cy5 and ApoB100-Cy3 are not transferred between lipoproteins and do not have altered clearance from plasma, or organ uptake (14–15).

### Confocal Analysis

The arterial retention of lipoproteins was quantified by a previously established method that measures the total fluorescent intensity obtained by confocal microscopy of at least five sections per carotid (Olympus IX-8 Perkin Elmer spinning disk microscope) (14–16). The volume of each section was calculated based on the area (determined by the elastin autofluorescence at 495 nm), the number of sections per carotid, and the interval between 10 Z-sections analyses per slice (as a volume) in order to minimize bias normally introduced within specific regions of interest. The total fluorescence intensity of the perfusate (532 nm for Cy3-LDL and 635 nm for Cy5-remnants) per unit volume was also collected for every experiment, under identical conditions in Z-series (in 3D as a volume). Based on the corresponding number of apoB48/100 particles (using a known concentration of ApoB48/100 in a specific volume of the perfusate) as well as the corresponding concentration of cholesterol per particle (14–16). The fluorescence quantitative analysis was carried out in a blinded fashion using ImageJ 1.50i software. Each carotid slide was assigned a distinctive code to ensure unbiased assessment.

### Solid-Phase Assays

#### Recognition to Chondroitin Sulfate and Extracellular Matrix

Recognition to CS was performed on amine-reactive 3D polymer-printed chamber slides (Nexterion Slide H, Schott Nexterion®, AZ, USA). The wells were incubated with 250 µL of the mAb solutions for 90 min at RT, diluted at 25 or 125 μg/mL in HBS containing 2 mmol/L CaCl_2_ and 2 mmol/L MgCl_2_ (sample buffer). Alternatively, the reactivity to extracellular matrix derived from VSMC (50,51) was evaluated in Lab Tek Chambered slides. In this case, the plates were blocked with a 1% BSA solution in HBS for 1h at RT, followed by incubation with 400 μL of the mAb solutions (5-125 μg/mL) in sample buffer for 1h at RT (37). In both sets of experiments, the mAb reactivity was detected by mean of a Cy3-conjugated goat anti-human IgG serum (Jackson Immunoresearch Laboratories, West Grove, PA), diluted 1:1000 in 0.1% BSA in HBS, and incubated for 90 mins at RT. Finally, the chamber separators were removed, and the slides were scanned at 532 nm (GenePix 4000B, San Jose, CA). Specificity was assessed using hR3 mAb as isotype-matched control.

#### Immunogenicity of chP3R99 mAb in Rats

The capacity of chP3R99 mAb (Ab1) to induce an idiotypic cascade of Abs in rats was evaluated by ELISA, based on a previous approach (52) (Figure 1). Briefly, the kinetics of induction of anti-idiotypic Abs (Ab2 response) and anti-anti-idiotypic Abs (Ab3 response, autologous anti-CS rat Abs), both in insulin resistant and control rats immunized with this mAb. All the assays were performed in triplicate for each condition and the coefficient of variation was <10%. Background values were less than 0.1.

#### Quantitative ELISA

The content of human IgG was quantitated both in the sera and aortic (tissue) homogenates from rats treated with chP3R99 or hR3 mAbs by a commercial ELISA kit (Sigma-Aldrich). Similarly, haptoglobin (ALPCO, NH, USA), aspartate aminotransferase and alanine transaminase (Elabscience, TX, USA), and β2-microglobulin (Novus Biologicals, CO, USA) were quantified in plasma collected at the ending time following the instructions provided by the manufacturers.

#### Immunoblot Analysis

Activated Immobilon-P PVDF membranes (Millipore Sigma, ON, Canada) spotted with decorin or biglycan (20 μg/well) were blocked with a 5% skim powder milk solution in 0.1% Tween 20-containing tris-buffered saline (TBST) [50 mM Tris HCl (pH 7.5), 150 mM NaCl]. The membranes were incubated ON at 4°C in agitation with the rat tissue homogenates from immunized rats at a protein concentration of 1.5 mg/mL in the blocking buffer. After washing three times with TBST for 15 min, membranes were incubated for 1 h at RT with HRP-conjugated anti-human IgG or anti-rat IgG secondary Abs (Jackson) diluted in blocking buffer. The reaction was visualized by chemiluminescence using ECL Advance Western Blotting Detection kit (GE HealthCare) and the dots quantified using ImageJ NIH software (version 1.50i).

#### Postprandial Studies and Lipid Analysis

Postprandial studies were conducted in rats at 15 weeks of age, 24 h after bolus injection of the mAbs (n=3 per group) or a week after the fifth immunization (n=6 per group). After fasting ON, rats were subjected to an oral fat challenge as described elsewhere (17). Blood samples were drawn from the tail at times 0, 2, 4, 5, 6 and 8 h following consumption of the pellet meal. Plasma triglycerides, total cholesterol, LDL cholesterol and high-density lipoproteins cholesterol were determined both in postprandial and fasting plasma samples by colorimetric assays (Wako, Chemicals USA Inc., Richmond, VA). The lipid content in liver lysates was quantified by high-performance liquid chromatography at the lipidomic core facility from the University of Alberta (Edmonton, AB, Canada), whereas triglycerides and total cholesterol were also measured by colorimetric assays in the homogenates (Wako). The lipids were expressed in terms of lipid mass per milligram of protein (17).

#### Cytokine Quantification

Cytokines, growth factors, leptin and the C-reactive protein were quantified by multiplex immunofluorescence assays by the company Eve Technologies (Calgary, AB, Canada) in aortic homogenates from rats and/or plasma.

#### Statistical Analysis

Statistical analysis was performed in GraphPad Prism 10.1.0 unless specified (GraphPad Software, Inc). Results are expressed as mean±SEM. Unpaired *t* test or Mann-Whitney *U* test (non-parametric data or n<6) were used to compare chP3R99-receiving groups with their corresponding isotype control-treated rats rather than comparing the acute treatment with the immunizations settings. For variables evaluated at different time points, a repeated measures two-way analysis of variance (ANOVA) followed by Tuckey post-hoc test was applied. Conversely, dose-response experiments were analyzed by ANOVA followed by a linear trend or Sidak post-tests. Other multiple comparisons were analyzed by a one-way ANOVA and Tuckey. For postprandial studies, the area under the curve (AUC) was determined in GraphPad Prism (17). Further bootstrapping analysis to compared postprandial AUC was performed in SPSS 25 for 1000 iterations to minimize bias and potential confounders. After the resampling, a Generalized Linear Model, not assuming equal variance, followed by the T3 Dunnett post-hoc test for multiple comparisons was applied. Values of *P*<0.05 were considered statistically significant. The significance levels are shown in the figures and the legends of the figures and tables.

## RESULTS

### ChP3R99 mAb Recognizes Vascular Extracellular Matrix Inhibiting Subendothelial Lipoprotein Retention in Insulin Resistant Rats

Solid-phase *in vitro* studies confirmed a specific capacity of chP3R99 mAb to recognize CS (Figure 2A, *P*=0.0143) and demonstrated a dose-dependent binding to the extracellular matrix secreted by VSMC (Figure 2B, *P*<0.0001). Previous studies have demonstrated that chP3R99 mAb competes with LDL for the binding site on sulfated GAG sugar chain of proteoglycans (47). Here we hypothesized that the chP3R99 mAb would interfere with the arterial retention of both LDL and remnant lipoproteins. Figure 2C-D shows that the binding of LDL to CS and the extracellular matrix *in vitro* was approximately 3.5-fold greater than for remnants. In this setting, chP3R99 significantly reduced the interaction of both LDL and remnants with either CS (Figure 2C, *P*<0.05) or the vascular extracellular matrix (Figure 2D, *P*<0.05). While the absolute reduction of the number of LDL particles resulting from the addition of chP3R99 mAb was greater than for remnants, the corresponding proportion of cholesterol-reduction was similar between the two lipoprotein preparations (Figure 2E-F, *P*<0.05). We also verified these results using VSMC cultured in hyperglycemic conditions in the presence of insulin (Figure S1A-C). Furthermore, the interaction of lipoproteins with CS was dose-dependent (*P*=0.0001) and highly sensitive to structural modifications of the particles. In our study, we employed copper-oxidized LDL as a representative model bearing a broad spectrum of oxidative modifications (Figure S1-D).

**Figure 2.**
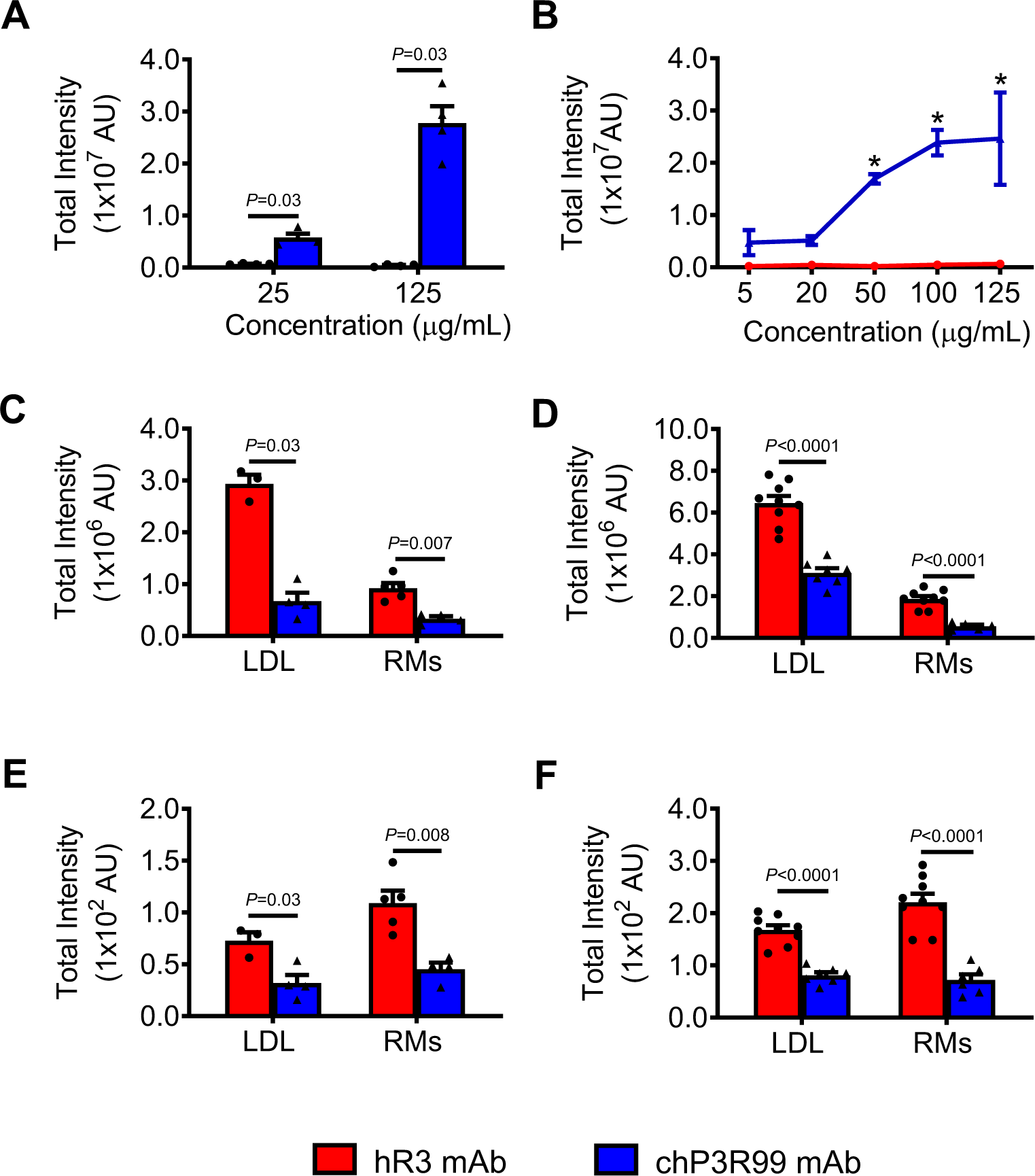
chP3R99 mAb blocking properties *in vitro*. Recognition of chP3R99 mAb to (A) Chondroitin sulfate (CS) and (B) extracellular matrix derived from vascular smooth muscle cells (VSMC) from rat aorta explants. Amino-reactive 3D chambered slides (Nexterion H) coated with CS (1 mg/mL) were incubated with chP3R99 mAb or the isotype-matched control (25 or 125 µg/mL) whereas extracellular matrix-coated Lab Tek II chambers were incubated with increasing concentrations of the mAbs (0-125 µg/mL). Recognition was detected with a Cy3-conjugated goat anti-human IgG serum. Inhibitory effect of chP3R99 mAb on lipoprotein binding to (A) CS and (B) extracellular matrix *in vitro*. Wells were pre-incubated for 90 min with the mAbs (250 μg/mL) followed by incubation with the lipoproteins (Cy3-LDL or Cy5-remnants) at 250 μg/mL. Slides were scanned at 520 nm and/or 640 nm to determine the fluorescence associated with specific binding (GenePix 4000B). Cholesterol associated with bound lipoproteins to CS (E) and the extracellular matrix (F) was estimated based on the total intensity and the concentration of cholesterol per particle of lipoprotein/intensity AU. Specificity was assessed using hR3 mAb as isotype-matched control. Results are expressed as the mean±SEM of at least three experiments. For unpaired two-group comparisons, Mann-Whitney *U* test or Student *t* test were used, \**P*<0.05. Two-way ANOVA followed by Sidak post-hoc test was applied to compare groups at different doses, *\*P*<0.001. RMs, remnants.

The findings *in vitro* prompted us to investigate whether chP3R99 could impinge the arterial retention of remnants under physiological conditions. As a first approach, carotid arteries from insulin resistant rats were pre-perfused (*in situ*) with chP3R99 mAb at a concentration of 125 µg/mL, followed by the perfusion of pre-labelled remnants at 90 µg/mL. Presence of chP3R99 significantly reduced the retention of both the number of remnant particles in arterial vessels (∼65%) as well as the corresponding deposition of remnant-associated cholesterol (Figure 3A, C, *P*<0.001). We also observed a dose-dependent capacity (25-250 µg/mL, *P*<0.0001) of the chP3R99 mAb to inhibit arterial retention of remnant lipoproteins *in vivo* (Figure 3D and Figure S2-A) as well as the corresponding deposition of cholesterol (Figure 3E, *P*<0.004). Unlike the isotype control mAb, abundant accumulation of chP3R99 *per se* was visualized in carotids from insulin resistant rats, in areas void of remnants (Figure 3F).

**Figure 3.**
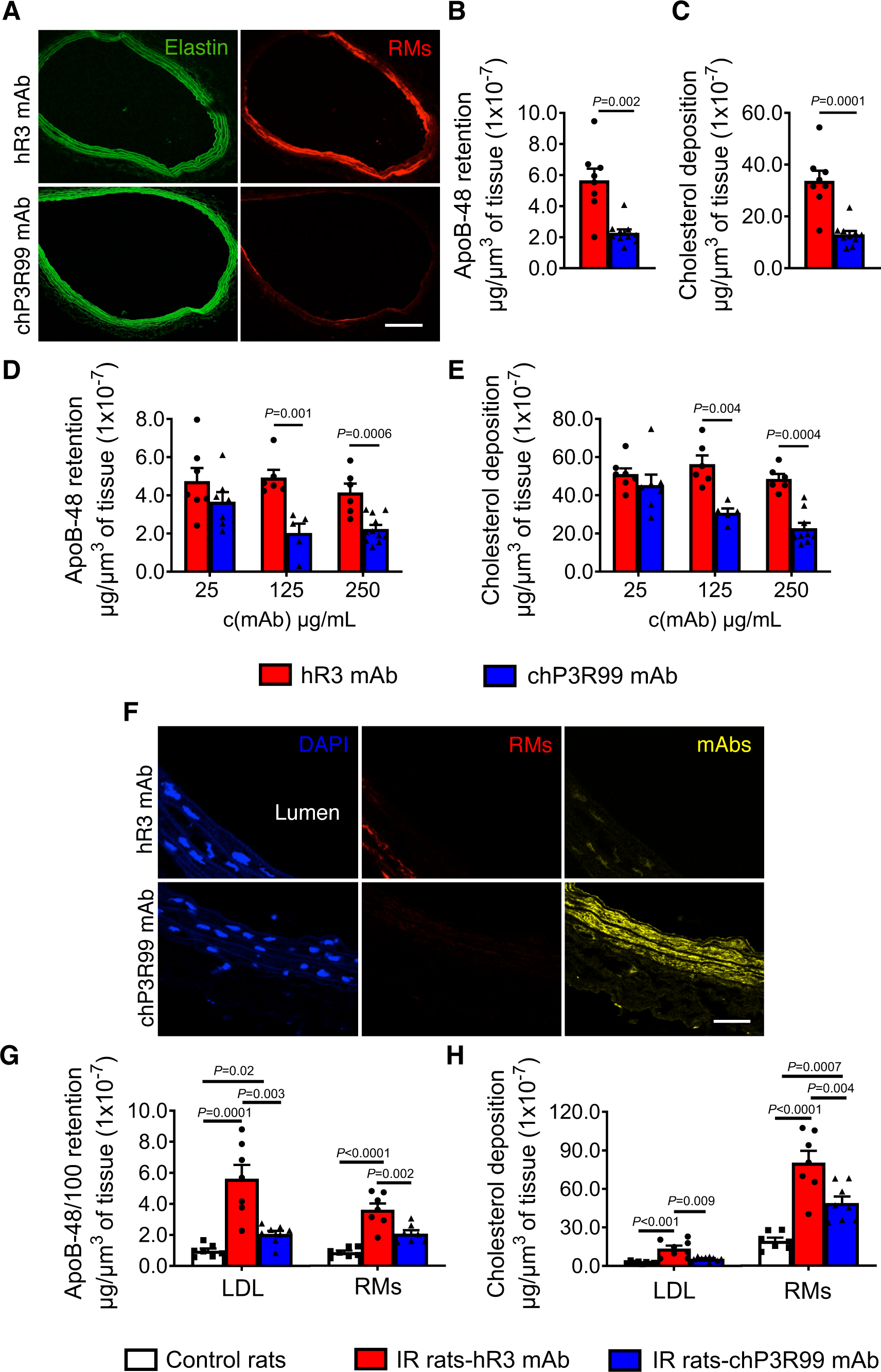
Acute effect of chP3R99 mAb on arterial retention of chylomicron remnants in insulin-resistant (JCR:LA-cp) rats. Carotid arteries from 12-week-old insulin resistant rats (n=4-6) were pre-perfused *ex vivo* with chP3R99 mAb for 20 min followed by the perfusion of fluorescently labeled lipoproteins. (A) Representative confocal images showing the blockade of Cy5-remnants (red) arterial retention (90 μg/mL) by the chP3R99 mAb (125 μg/mL). Morphology in green. Magnification 20X, Scale bar=100 µm. (B) Quantification of remnant particle number expressed as ApoB-48 molecules per volume unit. (C) Cholesterol deposition in carotids per volume unit. (D) Dose-response effect of chP3R99 on Cy5-remnants (150 μg/mL) arterial retention and (E) on cholesterol deposition in insulin resistant rats. (F) Representative images showing a reduction of Cy5-remnant retention (red) associated with the arterial accumulation of chP3R99 mAb perfused at 250 μg/mL (yellow). Counterstaining of nuclei in blue (DAPI). Magnification 100X, Scale bar=50 µm. (G) Relative inhibitory effect of chP3R99 on Cy5-remnant retention in the presence of equivalent concentrations of Cy3-LDL and (H) on cholesterol deposition. Total fluorescent intensity at 532 nm (LDL) and 635 nm (remnants) was determined and the corresponding number of ApoB-48/100 particles was estimated relative to the total intensity associated with a known concentration of ApoB in the perfusate, \**P*<0.05. Cholesterol content in the vessels was estimated as per ApoB, relative to the total intensity associated with cholesterol content in the perfusate. Lean rats were perfused solely with lipoprotein preparation as control whereas specificity was assessed by using the hR3 mAb as isotype-matched control. Data are mean ± SEM. For unpaired two-group comparisons, Mann-Whitney *U* test or Student *t* test were used. Dose-response effect was analyzed by ANOVA and linear trend post-test while multiple group comparisons were performed by one-way ANOVA followed by Tuckey, *P*<0.05.

Interestingly, *ex-vivo* (competitive) perfusion of equivalent amounts of both LDL and remnants in carotids from insulin resistant rats resulted in significantly higher retention of ApoB100 (3.6-fold) and ApoB48 (2.8-fold) respectively, relative to lean controls (Figure 3G). Pre-perfusion with chP3R99 reduced the retention of both LDL (∼65%) as well as remnants (∼30%). Despite the lower retention of the latter, its contribution to cholesterol deposition in the carotids was 6-fold greater than for LDL (80.4±9.1 x10^-7^ μg/μm^3^ *vs* 13.5±2.3 x10^-7^ μg/μm^3^, *P*<0.001) (Figure 3H). Also of note is that we employed a ratio of ApoB/P3R99 (90 µg/mL: 125 µg/mL respectively) to assess the blocking capacity of chP3R99 on remnant retention relative to LDL (non-saturating conditions in which the mAb elicit approximately 50% inhibition for LDL).

### Immunization with chP3R99 mAb Reduces the Arterial Retention of Pro-Atherogenic Lipoproteins in Insulin Resistant Rats

A unique property of the chP3R99 mAb is the immunogenicity of its idiotypic region (involved in antigen recognition), able to induce anti-sulfated GAG Abs *in vivo* (Figure 1) (40,42). Here we evaluated the immunogenicity of chP3R99 in JCR:LA-*cp* rats (Figure 4A), along with the impact of the idiotypic vaccination approach on lipoprotein retention. Immunization with chP3R99 mAb also resulted in a reduction of the retention of both LDL and remnants in the carotids of insulin resistant rats (Figure 4B). Quantitative analysis demonstrated a lower content of ApoB100 (∼30%, *P*=0.002) and ApoB48 (∼40%, *P*<0.0001) in the artery wall and the associated ApoB-cholesterol (Figure 4C-D, *P*<0.05).

**Figure 4.**
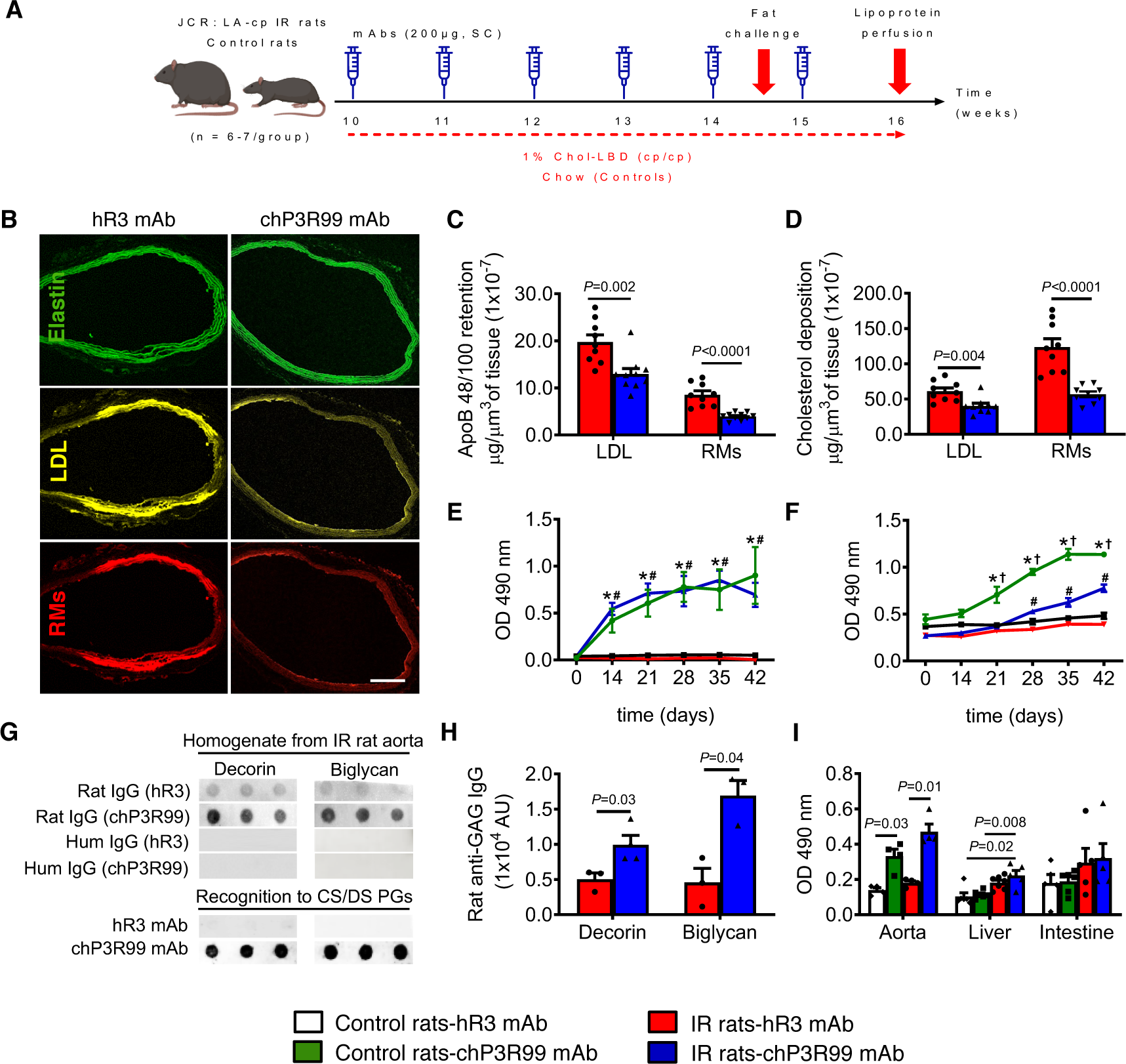
Effect of the vaccination with chP3R99 mAb on arterial lipoprotein retention in insulin-resistant (IR) rats (JCR:LA-cp). (A) Study Design: Both insulin resistant and control rats (n=6-7) received six subcutaneous injections of chP3R99 mAb or the isotype-matched control hR3 (200 µg/week). Vaccinated insulin resistant rats underwent an *ex-vivo* carotid perfusion of equivalent amounts of Cy3-LDL and Cy5-remnant lipoproteins (150 µg/mL). (B) Representative images of the arterial retention of remnants (red) and LDL (yellow) in insulin resistant rats. Morphology in green. Magnification 20X, Scale bar=100 µm. (C) Quantification of lipoprotein retention in the vessels (n=5 sections/animal, 10-Z sections/slice). The corresponding number of ApoB-48/100 particles was estimated relative to the total intensity associated with a known concentration of ApoB in the perfusate, *P*<0.05. (D) Arterial cholesterol deposition relative to the total intensity associated with a known concentration of cholesterol in the perfusate, *P*<0.05. (E) Kinetics of induction of anti-idiotypic Abs (Ab2) in rats immunized with chP3R99 mAb. The Ab2 response was assessed by subtracting the reactivity of the sera to the isotype-match control (anti-isotypic response) from the total response directed to chP3R99 by ELISA. (F) Kinetics of induction of anti-CS Abs in rats (Ab3 response). (G) Detection of Ab3 within the aorta by Dot Blot. Membranes were spotted with CS-PGs (20 µg/dot) and incubated with aorta homogenates (1 mg/mL) from immunized rats. (H) Relative quantification of anti-CS Abs in aorta homogenates. (I) Accumulation of anti-CS Abs in tissue lysates by ELISA. GAG-coated plates (10 µg/mL) were incubated with rat sera (1/200) or the tissue lysates (1 mg/mL). The presence of rat IgG in the tissues was detected by an anti-rat IgG secondary serum whereas the recognition of P3R99 to proteoglycans or its presence in the aorta homogenates were assessed utilizing an anti-human IgG secondary serum conjugated to HRP. Data are mean±SEM. For unpaired two-group comparisons, Mann-Whitney *U* test or Student *t* test were used, *P*<0.05. Two-way ANOVA followed by Tuckey post-hoc test was applied for repeated measures. *†*P*<0.001, chP3R99-immunized groups compared with their corresponding isotype-match controls. #*P*<0.001, chP3R99-immunized IR rats compared with chP3R99-immunized control rats. Multiple group comparisons were performed by Kruskal-Wallis test followed by Dunn’s post-hoc test, *P*<0.05.

Kinetic experiments revealed that chP3R99 mAb (but not the isotype control), mounted a strong anti-idiotypic (Ab2) response in rats after a second injection of the mAb, which was very similar between wildtype and insulin resistant rats (Figure 4E, *P*>0.05). Interestingly, significantly higher anti-CS Abs (Ab3 response) were detected in sera three weeks after the first mAb injection in the lean controls (*P*<0.0001) while requiring an additional week for insulin resistant rats (*P*=0.0003) when compared with the isotype control. Overall, the levels of anti-CS Abs induced by the chP3R99 mAb were higher in the sera of lean rats when compared with insulin resistant rats after three weeks of immunizations (Figure 4F, *P*<0.0001). We also found an induction of anti-dermatan sulfate Abs in contrast to the unsulfated GAG hyaluronic acid in rats (Figure S3-A).

Since the immunogenicity of chP3R99 appeared similar in lean and insulin resistant rats, we hypothesized that lower circulating levels of anti-CS Abs in the later could be related to the accumulation in the arteries rather than an impaired induction in insulin resistant rats. To address this, we developed an immunoblot technique in which both decorin and biglycan were spotted on membranes (as examples of proatherogenic CS/dermatan sulfate-containing proteoglycans), followed by incubation with aorta homogenates. We detected the presence of autologous anti-proteoglycans IgG (Ab3) generated by the immunization in aortic lysates from insulin resistant rats immunized with chP3R99 (Figure 4G-H), which was further confirmed by ELISA (Figure 4I). Neither this mAb (Figure 3B), nor the autologous anti-CS Abs significantly accumulated in organs relevant for lipoprotein metabolism like the liver and the intestine (Figure 4I). Conversely, fluorescent studies did not show significant accumulation of chP3R99 in aorta due to the low dose of this mAb employed for immunization (Figure S3).

### Treatment with chP3R99 Did Not Affect the Overall Biochemical Status of Rats

Additional studies were conducted in lean and insulin resistant rats to address potential toxicity associated with a therapy directed to CS, not only in normal conditions but also in a more compromised physiological status of insulin resistance. We analyzed clinical and biochemical parameters in rats immunized with chP3R99 (Figure 4A) or those who received a bolus IV injection of this mAb (Figure 5A).

**Figure 5.**
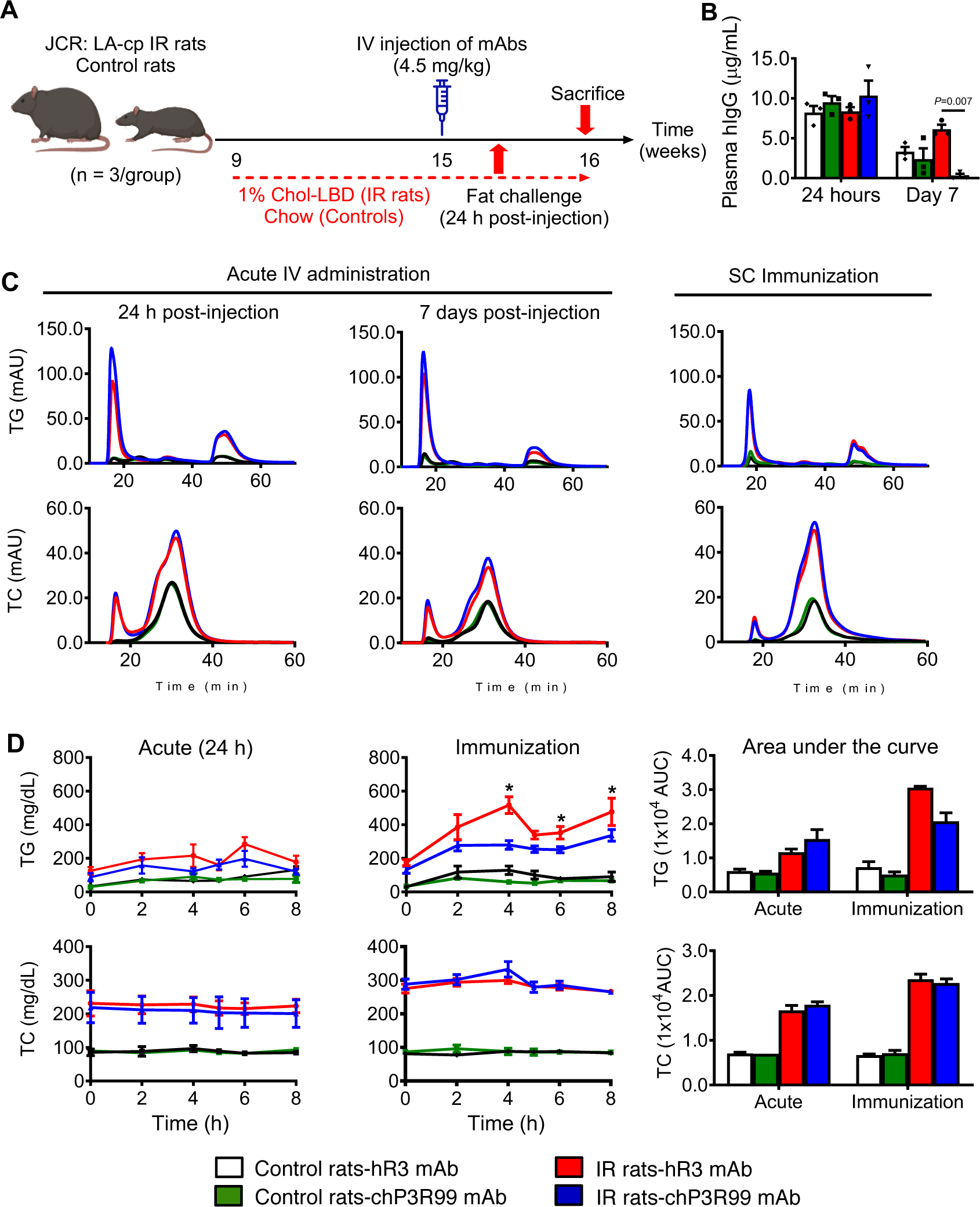
Effect of acute administration of chP3R99 mAb and the immunization with this mAb in lipoprotein metabolisms. (A) Acute intravenous administration of chP3R99 mAb in lean control and cholesterol-fed insulin resistant (IR) rats (n=3). The rats were inoculated with 4.5 mg/kg (body weight) of chP3R99 mAb in PBS through the jugular vein at 15 weeks of age. (B) Quantification of human IgG in rat sera 24 hours and seven days after bolus administration of the mAbs. (C) Triglyceride (TG) and total cholesterol (TC) FPLC fractions in pooled plasma from rats subjected to intravenous inoculation of or immunization with chP3R99 mAb. (D) Postprandial study in rats. At 15 weeks of age, the rats were fasted overnight and subjected to an oral fat challenge. Specificity was assessed using hR3 mAb as isotype-matched control. Results are expressed as mean±SEM. Multiple comparisons were performed with Kruskal-Wallis followed by Dunn’s post-hoc test or by a repeated measure two-way ANOVA followed by Tuckey, *P*<0.05.

Primarily, we assessed the serum levels of human IgG in each experimental condition to assess the direct impact of chP3R99 mAb on the overall physiological state of the rats. In the immunization experiment, quantitative ELISA only revealed detectable circulating levels of the isotype control mAb both in lean controls (104.6±21.8 ng/mL) and insulin resistant rats (63.1±39.7 ng/mL) but not of chP3R99 mAb. Although all groups exhibited comparable levels of circulating IgG after 24 h post injection in the acute setting, insulin resistant rats administered chP3R99 also displayed significantly reduced concentrations after one week (Figure 5B, *P*<0.001). Consistent with this finding, we specifically demonstrated the accumulation of chP3R99 mAb in aorta homogenates from insulin resistant rats (10.6±1.0 ng/mg of protein) while dot blot studies did not show any accumulation in the liver or intestine (Figure S3B).

As expected, 16-week-old cholesterol-fed JCR:LA-cp rats were characterized by significantly higher levels of leptin (153.1±13.8 vs 34.7±2.6 µg/mL, *P*<0.0001), obesity (584.5±10.2 vs 337.5±10.1 g, *p*<0.0001) and dyslipidemia (Table 1 and Figure 5), relative to age-matched control rats (Table S1). Importantly, the body weight was not affected throughout the treatments. Although no significant fasting hyperglycemia was observed (*P*>0.05), extremely high levels of insulin were detected (1228.0±84.3 vs 129.5±10.6 pmol/L, *P*<0.0001), confirming severe insulin resistance as indicated also by their HOMA-IR index (Table 1). Likewise, insulin resistant rats showed significantly higher levels of aspartate aminotransferase and alanine aminotransferase than the lean controls (*P*<0.01), measured as indicators of liver damage. Plasma concentration of β2-microglobulin, a surrogate marker of kidney damage, was also significantly increased in insulin resistant rats (*P*<0.01). None of the parameters mentioned above were affected by the treatment with chP3R99 (Table 1 and Table S1). Neither the acute treatment with chP3R99 nor the immunization with chP3R99 affected the total blood cells counts in insulin resistant or lean control rats (Tables S2 and S3).

**Table 1.** Biochemical and Clinical Parameters in Insulin Resistant Rats Treated with chP3R99 mAb.

| Parameter | Acute injections (4.5 mg/kg, IV) |  | Immunizations (200 µg, SC, 6 wk) |  |
| --- | --- | --- | --- | --- |
|  | hR3 mAb | chP3R99 mAb | hR3 mAb | chP3R99 mAb |
|  | (n=3) | (n=3) | (n=5) | (n=6) |
| Glucose, mmol/L | 7.9±0.5 | 8.0±0.8 | 11.5±1.5 | 9.6±0.9 |
| Insulin, pmol/L | 1058.0±170.3 | 1270.0±189.4 | 1260±195.0 | 1267 ±134.6 |
| HOMA-IR, AU | 147.5±21.3 | 169.1±8.3 | 69.6±8.1 | 73.3±7.6 |
| Leptin, µg/mL | 56.9±11.2 | 65.7±13.5 | 119.7±13.4 | 101.5±15.1 |
| Plasma LDL, mmol/L | 2.0±0.2 | 1.6±0.1 | 1.8±0.1 | 2.8±0.1 |
| Plasma cholesterol, mmol/L | 5.4±0.4 | 4.6±0.2 | 7.4±0.4 | 7.1±0.3 |
| Plasma triglycerides, mmol/L | 3.7±0.5 | 4.9±0.4 | 2.0±0.2 | 1.9±0.2 |
| Liver cholesterol, µg/mgPr | 36.7±4.8 | 32.7±2.3 | 30.6 ±2.4 | 31.3±5.3 |
| Liver triglycerides, µg/mgPr | 143.5±15.8 | 162.4±15.9 | 96.8±13.9 | 112.4±7.3 |
| Aspartate aminotransferase, µg/mL | 20.8±1.5 | 20.2±0.4 | 20.7±0.9 | 19.6±2.1 |
| Alanine aminotransferase, µg/mL | 6.0±0.9 | 6.2±1.4 | 7.1±1.0 | 5.1±2.2 |
| Haptoglobin, mg/L | 11.3±2.7 | 3.4±1.6* | 10.6±3.1 | 9.6±1.9 |
| C reactive protein, mg/L | 231.7±44.8 | 300.2±31.9 | 311.9±55.4 | 350.3±37.7 |
| β2-Microglobuline, µg/mL | 7.8±2.4 | 7.6±2.0 | 8.0±1.2 | 9.5±1.3 |

Results are mean±SEM. Two-tailed Mann-Whitney *U* test was applied to compare chP3R99-receiving groups with their corresponding isotype-matched controls. \**P*<0.05, †*P*<0.01. HOMA-IR: Homeostatic Model Assessment for Insulin Resistance.

### ChP3R99 mAb Did Not Interfere with Lipoprotein Metabolism *In Vivo*

The role of heparan sulfate proteoglycans, particularly syndecan I, in the hepatic clearance of triglyceride-rich lipoprotein remnants has been well established (53). To study in more detail whether a treatment based on chP3R99 mAb could affect lipoprotein metabolism we analyzed plasma triglycerides and cholesterol levels at different time points, both in fasting conditions and in the postprandial state (Figure 5, Table 1).

In fasting samples (Figure 5C), insulin resistant rats under high-fat diet displayed higher levels of plasma triglycerides (mainly associated with the chylomicron/very low-density lipoproteins and high-density lipoproteins fractions) relative to the lean controls. Total cholesterol was also elevated in those rats but mostly distributed in high-density lipoproteins and less extensive in the chylomicron/very low-density lipoproteins fraction. Importantly, the bolus administration of chP3R99 mAb did not modify the lipid profile of insulin resistant or lean rats when compared with the isotype-matched control 24 h or a week after the mAb administration. Similar results were observed in the rats immunized with chP3R99 mAb. Further assessment of triglycerides and cholesterol associated with different lipoprotein classes confirmed no difference between the groups treated with chP3R99 and their corresponding control groups (Table 1 and Table S1, *P*>0.05).

Assessment under the postprandial state (Figure 5D and Figure S4) demonstrated that the IV inoculation of chP3R99 had no impact on plasma concentrations of triglycerides, total cholesterol, and LDL. Interestingly however, insulin resistant rats immunized with chP3R99 showed significantly lower triglyceride levels at 4h (*P*<0.0001), 6h (*P*=0.021) 8h (*P*=0.014) after a fat challenge, relative to hR3-receiving rats (Figure 5D). Further bootstrapping analysis of the AUC corresponding to postprandial triglycerides (Figure 5D) confirmed that immunization of insulin resistant rats with the chP3R99 mAb improved triglyceride metabolism relative to the ones immunized with the isotype-matched control [AUC delta = 888.9] [CI BCa 95%, -1625.9 to -142.5], while no differences were detected in the control rats [AUC delta = 262.9] [CI BCa 95%, -55.7 to 592.5].

Histological analysis of liver sections depicted steatosis and fibrosis in insulin resistant rats (Figure 6A) while the treatment with chP3R99 did not impact the hepatic neutral lipid content (Figure 6B). Further, colorimetric assays (Table 1 and Table S1) and high-performance liquid chromatography (Tables S4 and S5) in liver lysates confirmed no modulation of hepatic lipids by chP3R99 in any of the groups.

**Figure 6.**
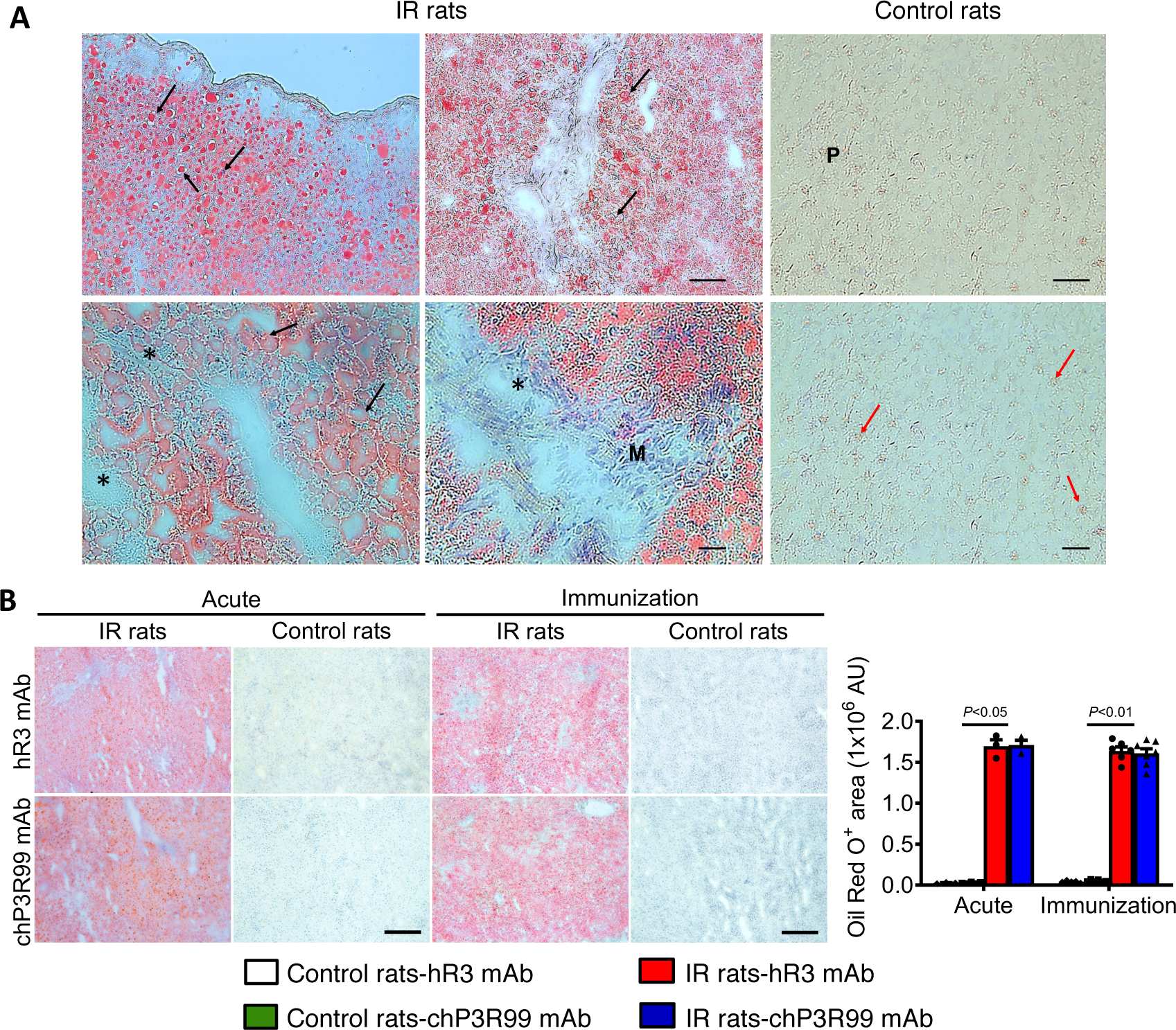
Effect of acute administration of chP3R99 mAb or the immunization with this mAb in hepatic neutral lipids content. (A) Histopathological findings in liver sections from insulin resistant (IR) rats fed a 1% cholesterol-enriched diet versus control rats. Oil Red O staining revealed steatosis in IR rats (red) characterized by a high content of extracellular and intracellular lipid vacuoles (black arrows), fibrosis (asterisks), and mononuclear cell infiltration in some regions (M). Control rat showed a normal parenchyma (P) and scarce lipid content associated to hepatocytes (red arrow). Magnification 20X (top panel A, bar=150 µm) and magnification 40X (bottom panel A, bar=50 µm). (B) Neutral lipids content in the liver of rats visualized by oil red O staining. Scale bar=100 μm; Magnification 20X. Five images were quantified per animal. Specificity was assessed using hR3 mAb as isotype-matched control. Results are expressed as mean±SEM. Multiple comparisons were performed with Kruskal-Wallis test followed by Dunn’s post-hoc test, *P*<0.05.

### ChP3R99 mAb Did Not Affect Plasma Concentrations of Inflammatory Cytokines in Insulin Resistant Rats

Unlike the C-reactive protein (which was similar in all the groups), plasma levels of haptoglobin were significantly higher in insulin resistant rats compared to lean controls (10.9±1.9 vs 3.6±0.5 mg/mL, *P*=0.006). Interestingly, a 3-fold reduction in the concentrations of haptoglobin was observed in insulin resistant rats who received an acute administration of chP3R99 (Table 1, *P*<0.05), to similar levels as the lean controls, while showing no effect in the lean control group or any of the immunized rats (Table 1 and Table S1). As expected, we verified a low grade systemic inflammatory condition in insulin resistant rats relative to control (Table 2); characterized by increased levels of interleukin-6 (*P*<0.0001) and of interferon-γ, tumor necrosis factor-α, interleukin-2, granulocyte colony-stimulating factor and the C-C motif chemokine ligand-5 (CCL5/RANTES) (*p*<0.05). However, no significant difference was observed between any of the mAb-treated groups in insulin resistant rats treated with chP3R99 mAb when compared with their corresponding isotype control-treated counterparts (*P*>0.05).

**Table 2.** Cytokine and Growth Factors Levels in Plasma from Insulin Resistant Rats Treated with chP3R99 mAb.

| Parameter | 10-wk-old rats |  | Acute mAb injections in IR rats<br>(4.5 mg/kg, IV) |  | Immunizations in IR rats<br>(200 µg, SC, 6 wk) |  |
| --- | --- | --- | --- | --- | --- | --- |
|  | Control rats | IR rats | hR3 mAb | chP3R99 mAb | hR3 mAb | chP3R99 mAb |
|  | (n=5) | (n=5) | (n=3) | (n=3) | (n=5) | (n=6) |
| G-CSF, pg/mL | 112.4±17.1 | 230.1±30.6† | 238.1±34.8 | 168.2±19.5 | 200.4±37.0 | 132.8±32.1 |
| GM-CSF, pg/mL | 130.4±12.9 | 306.5±169.3 | 426.1±286.1 | 351±112.2 | 646.4±148.6 | 543.0±239.6 |
| Eotaxin, pg/mL | 13.5±2.5 | 18.6±2.1 | 20.1±1.3 | 12.2±1.9 | 25.6±4.5 | 39.7±7.6 |
| MIP-1a, pg/mL | 21.9±1.6 | 32.4±4.4 | 23.9±9.5 | 18.9±3.5 | 24.7±6.6 | 38.4±7.8 |
| MIP-2, pg/mL | 309.7±11.8 | 357.0±35.3 | 405.4±61.4 | 311.2±34.6 | 439.3±52.2 | 465.0±62.8 |
| MCP-1, µg/mL | 1.2±0.1 | 1.5±0.2 | 2.1±0.7 | 1.1±0.3 | 2.4±0.6 | 2.3±0.7 |
| Fractalkine, pg/mL | 84.7±2.2 | 114.3±14.6 | 81.4±18.3 | 80.9±26.9 | 73.6±15.6 | 147.4±50.5 |
| CCL5, pg/mL | 1.9±0.3 | 3.2±0.4* | 0.3±0.0 | 0.3±0.0 | 1.1±0.5 | 1.6±0.4 |
| LIX, µg/mL | 3.7±0.7 | 4.8±0.9 | 0.6±0.1 | 0.5±0.1 | 3.8±0.8 | 2.7±1.1 |
| IP-10, pg/mL | 594.1±17.5 | 663.3±91.3 | 475.4±67.4 | 431.4±56.6 | 516.3±80.7 | 511.3±49.3 |
| CXCL1, pg/mL | 503.4±129.4 | 721.1±113.7 | 556.3±247.3 | 214±25.2 | 566±198.5 | 809.2±196.1 |
| EGF, pg/mL | 10.9±5.4 | 8.4±2.8 | 4.3±4.1 | ND | 19.1±12.9 | 22.4±13.2 |
| VEGF, pg/mL | 33.8±15.6 | 95.6±32.6 | 136.7±54.2 | 54.9±45.4 | 64.6±35.6 | 126.9±77.2 |
| TGF-β1, µg/mL | 27.4±12.5 | 40.9±10.5 | 1.8±1.6 | 1.2±0.6 | 34.7±1.1 | 23.3±12.7 |
| TGF-β2, µg/mL | 1.4±0.6 | 1.8±0.4 | 0.2±0.0 | 0.1±0.0 | 1632.0±573.8 | 877.9±454.7 |
| IFN-γ, pg/mL | 343.8±248.0 | 1097.0±189.4* | 778.4±638.1 | ND | ND | ND |
| TNF-α, pg/mL | 16.5±2.5 | 32.5±7.2* | 87.1±53.5 | 37.8±4.1 | 49.2±12.8 | 61.7±23.9 |
| IL-1α, µg/mL | 198.5±26.9 | 437.3±115.5 | 199.5±143.6 | 54.9±23.6 | 278.4±84.8 | 129.0±15.7 |
| IL-1β, pg/mL | 1429.0±461.1 | 947.2±289.6 | 2221.0±450.9 | 977.4±357.8 | 3800.0±1820.0 | 2935.0±623.5 |
| IL-2, pg/mL | 77.4±10.6 | 161.1±32.1* | 204.4±112.7 | 152.4±90.9 | 250.8±83.6 | 262.0±35.4 |
| IL-5, pg/mL | 99.8±2.9 | 123.9±11.4 | 179.4±74.5 | 90.8±10.8 | 162.9±22.9 | 236.3±46.5 |
| IL-6, µg/mL | 34.7±2.6 | 153.1±13.9‡ | 147.5±21.3 | 169.1±8.3 | 119.7±13.5 | 101.5±15.1 |
| IL-10, pg/mL | 1242.0±422.9 | 839.9±242.2 | 2084.0±487.9 | 913.3±332.7 | 3304.0±1567.0 | 2555.0±497.0 |
| IL-12p70, pg/mL | 222.5±48.3 | 531.2±172.0 | 1648.0±1037.0 | 176.9±39.6 | 668.7±178.3 | 627.1±183.1 |
| IL-13, pg/mL | 17.4±5.8 | 29.9±13.2 | 104.1±72.4 | 1.46±1.0 | 64.4±15.1 | 59.9±21.9 |
| IL-17A, pg/mL | 25.5±7.9 | 42.9±17.4 | 110.6±70.3 | 15.9±4.6 | 62.4±20.9 | 44.6±14.5 |
| IL-18, pg/mL) | 292.9±71.7 | 518.3±140.9 | 295.7±85.1 | 325.9±144.1 | 271.8±83.9 | 608.2±330.2 |

Results are mean±SEM. Two-tailed Mann-Whitney *U* test to compare chP3R99-receiving groups with their corresponding isotype-match controls or chow-fed insulin resistant rats versus the lean control. \**P*<0.05, †*P*<0.01, ‡*P*<0.01. ND, not detected; G-CSF, Granulocyte colony-stimulating factor; GM-CSF, Granulocyte macrophage colony stimulating factor; MIP, Macrophage inflammatory protein; MCP, Monocyte chemoattractant protein; CCL5 (RANTES), C-C motif chemokine ligand 5; LIX, Lipopolysaccharide-inducible CXC chemokine; IP-10, Interferon gamma-induced protein 10; EGF, Epidermal growth factor; CXCL1 (GRO/KC), Chemokine (C-X-C motif) ligand 1; VEGF, Vascular endothelial growth factor; TGF, Transforming growth factor; IFN-γ, Interferon gamma; TNF-α, Tumor necrosis factor alpha; IL, Interleukin.

A similar cytokine array performed in aorta homogenates in insulin resistant rats showed no differences between the groups (*P*>0.05) for the variables reliably detected in those samples (Table S6).

## DISCUSSION

Epidemiological and Mendellian Randomization studies in Europe, North America and now other countries (9–13), support the notion that remnant lipoproteins (measured as remnant cholesterol), have substantial independent contributions to CVD risk; and that these relationships may be stronger than for LDL-C under conditions of residual risk and/or diabetes. There is certainly a plethora of pre-clinical evidence demonstrating the interaction of both LDL and remnant lipoproteins within the arterial wall (3, 4, 8, 14–20), consistent with the response-to-retention hypothesis and attachment via select CS arterial proteoglycans (3, 4, 8, 20–23). Limited case-control studies have shown similar observations in human vascular tissue (20, 21, 54, 55). In order to advance our understanding of not only the contribution of these atherogenic lipoproteins, but to develop a potential therapeutic to interfere directly with the deposition of cholesterol, we sought to assess the efficacy of the chP3R99 mAb both *in vitro* and *in vivo* using a rat model of insulin resistance.

### Idiotypic Characteristics of chP3R99 mAb

The chP3R99 has been developed to recognize sulfated GAGs found within the chondroitin sulfate family of proteoglycans (37–39, 56, 57). In addition to antigen recognition mediated by the complementarity-determining region (CDR) motifs of the P3 idiotype, the region contains both CD4^+^ and CD8^+^ T-cells epitopes, conferring a high immunogenicity which is preserved within the chP3R99 mAb. The idiotope is a generic ‘structural’ term given to unique sequence of antigenic determinants (epitopes) located on the variable portion of any given Ab. An ‘anti-idiotype’ Ab [2^nd^ generation-Ab2], in turn, is specific for an Ab1 idiotope and may resemble the 3D structure of the antigen (Ab2β, also known as ‘internal image’ of the antigen) as described in the cascading ‘immune network theory’ originally proposed by Nobel Laureate Niels Jerne (reviewed in 48). The ‘anti-idiotypic response’ is thought to explain the amplification of a highly specific region of the ‘original’ idiotype and acts as a vaccine to provide long-term recognition (Figure 1B).

### ChP3R99 mAb Competes with Lipoproteins and Mounts an Anti-idiotypic Response

The therapeutic potential of the chP3R99 idiotypic mAb is two-fold; (i) acute recognition to the antigen, specifically its idiotype (in this case interfering with ApoB-GAG-complex formation), and (ii) vaccination with the mAb (chronic immunizations) mounts an autologous ‘cascade’ of Abs that reproduce the specificity of the original mAb, amplifying antigen recognition analogous to a vaccine. In this study we not only show that the chP3R99 mAb recognizes vascular extracellular matrix inhibiting subendothelial lipoprotein retention (acutely) in insulin resistant rats, but also that the ‘vaccination’ with the same, also can interfere with arterial retention of pro-atherogenic lipoproteins over time. Indeed, we have also been able to demonstrate the immunogenicity of chP3R99 inoculation in rats to develop not only Ab2, but Ab3 antibodies. Our kinetic experiments also revealed that the chP3R99 mAb mounted a strong anti-idiotypic (Ab2) response in rats after a second injection of the mAb, (similar in wildtype and insulin resistant rats). Interestingly, the detection of an Ab3 response in sera was delayed compared to the appearance of Ab2, indicating that the latter may be essential to induce an anti-CS response or activate/amplify a natural response against this antigen. Perhaps more interesting, was that the anti-CS Abs (Ab3 response) in insulin resistant rats was one week slower than for lean control wildtype rats; and overall, Ab levels did not reach the maximum response observed in control rats. It might be plausible that under systemic low grade inflammatory conditions the strength of the idiotypic response to chP3R99 may be impaired. However, the presence of a comparable Ab2 response, in both immunocompromised and normal conditions, suggests that the anti-CS Abs generated in insulin resistant rats could be accumulating in the arteries. Indeed, we were able to demonstrate, for the first time, that the autologous anti-CS Abs accumulate in aorta and associated with reduced lipoprotein retention *in vivo*. Other unpublished data from our group suggests that as few as two SC injections of chP3R99 (50 µg) were sufficient to interfere (reduce) aortic lipoprotein retention in Sprague-Dawley rats following a bolus administration of LDL. To our knowledge, the specific accumulation of Ab3 in the targeted organ has not been previously reported (as opposed to the detection Ab3 in sera or characterized *in vitro*) (58).

The clinical approach to the idiotypic theory has primarily relied on anti-idiotypic vaccines (Ab2 mAbs) (58), whereas chP3R99 mAb is an Ab1 capable of mounting a full idiotypic cascade. This unique characteristic not only enables chP3R99 to directly recognize CS-proteoglycans, but also opens the potential for its use as a passive therapy, a vaccine, or a combination of both strategies. Thus, the potential of this mAb in clinical applications extends beyond the direct antigen recognition amplifying its specificity through an idiotypic cascade mechanism.

While idiotypic/anti-idiotypic vaccines hold potential for targeting various antigens, a significant portion of the clinical applications explored so far have focused on carbohydrates and glycolipids, primarily in the field of cancer immunotherapy (summarized in 58). While carbohydrates are known for their poor immunogenicity, the fact that Ab2 resemble the antigens offer a promising strategy for mimicking those molecules (ie: GAG) within a protein framework, thereby potentially eliciting a more effective immune response. Future studies should now consider larger and higher order animal models to ensure that chronic inoculations of chP3R99 are sufficiently immunogenic in clinically translatable immune systems.

### ChP3R99 mAb Interactions with Specific Lipoproteins

Another interesting insight came from comparing the impact of single lipoprotein retention experiments (Figure 3), with double labelled lipoproteins. When both lipoproteins are compared, it is evident that the effect of chP3R99 is greatest on LDL by particle number (ie when measured as ApoB100). In contrast, the chP3R99 had a much larger effect by proportion on remnant lipoproteins when expressed as lipoprotein-associated cholesterol. These observations raise important factors regarding the size of the different particles (27 nm for LDL and ∼50 nm for remnants), the greater rate of permeability for LDL (due to the smaller size), yet greater amounts of proportional cholesterol for remnants (due to their larger volume). It is plausible that the chP3R99 mAb may indeed compete for these binding sites equally but impact lipoproteins differentially due to these factors. Additional follow up studies such as testing an upper limit of efficacy for these interactions *in vivo* maybe required. Another important note is that the current studies have been conducted with native forms of both LDL and remnants. We did test the binding of copper oxidized LDL versus native forms to chondroitin sulfate *in vitro* (Figure S1D) which suggested a reduced interaction with particles following modification. We also recognize that clinically, remnant cholesterol includes cholesterol from a heterogenous population of particles (of differing sizes), which is not completely reflected in the homogenous remnant preparations used in these series of experiments.

### Developmental Limitations of chP3R99 mAb

The development of a putative mAb to target vascular lipoprotein interactions does not come without theoretical limitations. The biological purpose of CS-proteoglycans is not limited to structural integrity of the vasculature. The biological function of the four main proteoglycan families is known to be expressed throughout physiology in extracellular, pericellular, cell surface and intracellular regions (59). The putative interaction of a mAb to each or any of these ‘on-target but off-vasculature’ sites would claim additional properties to impair functional (and/or structural) biochemical processes (hepatic HS-proteoglycans), compete with normal metabolic pathways (lipid metabolism) and plausibly stimulate inflammatory cascades. However, it is also notable that GAG composition in peripheral tissues is often modified in numerous pathologies, including ACVD and cancer. In atherosclerosis, CS-proteoglycans are predominantly synthesized by VSMCs, resulting in distinctive features such as enlarged GAG chains, increased sulfation, along with changes in the sulfation pattern (60).

In this study (using rats), we have not been able to identify any significant biochemical signal associated with any negative attribute of the chP3R99 mAb. Indeed, all studies that have utilized the chP3R99 mAb to date, have not reported any systemic ‘on-target’ but ‘off-vasculature’ effects including those in mice, rats and rabbits. To the contrary, there has been enticing data collected using immunoscintigraphy by intravenous administration of ^99m^Tc-chP3R99 mAb in atherosclerotic and control rabbits (47). The overall uptake of ^99m^Tc-chP3R99 mAb was predominantly localized in liver, kidneys and heart alongside all time intervals. However, the planar images acquisition revealed the accumulation of ^99m^Tc-chP3R99 mAb in the carotid of atherosclerotic rabbits 6 h after radiotracer administration, but not in control animals. Surprisingly, the immunization with this mAb showed an improvement in systemic postprandial triglycerides levels during insulin resistance without affecting the hepatic lipid content. This finding requires further attention to identify whether the induction of anti-sulfated GAG antibodies could indeed improve the endothelial chylomicron metabolism mediated by lipoprotein lipase, a process in which the role of proteoglycans has been described (61). Future studies will need to continue to explore various and reproducible ways of demonstrating inert clearance mechanisms of idiotype-derived mAbs *in vivo*.

## CONCLUSION

Acute and long-term immunization of chP3R99 mAb interfered with the interaction of both LDL and remnant lipoproteins with arterial proteoglycans in a setting of insulin resistance. These *in vivo* data support the innovative approach of targeting pro-atherogenic lipoprotein retention by chP3R99 as a passive therapy or as an idiotypic vaccine for atherosclerosis.

## ACKNOWLEDGEMENTS

We are grateful for the technical expertise provided by Ms. Sharon Sokolik.

## NONSTANDARD ABBREVIATIONS AND ACRONYMS

Ab: antibodies
Ab2: anti-idiotypic antibodies
Ab3: anti-anti-idiotypic antibodies
ASVD: atherosclerotic vascular disease
CS: chondroitin sulfate
DS: dermatan sulfate
ECM: extracellular matrix
GAG: glycosaminoglycan
IR: insulin resistant
LDL: low-density lipoprotein
mAb: monoclonal antibody
PG: proteoglycan
RM: remnants
VSMC: vascular smooth muscle cells

