## Supplemental materials for "Monoclonal antibody chP3R99 reduces subendothelial retention of atherogenic lipoproteins in Insulin-Resistant rats: *Acute treatment versus long-term protection as an idiotypic vaccine for atherosclerosis*"

**Running Title:** Atheroprotective mAb in insulin-resistant rats

**Authors:** Yosdel Soto^1,2*^, Arletty Hernández^1^, Roger Sarduy^1^, Victor Brito^1^, Sylvie Marleau^3^, Donna F. Vine^2^, Ana M. Vázquez^4^ and Spencer D. Proctor^2*^

**Affiliations:**
^1^ Department of Immunobiology, Direction of Immunology and Immunotherapy, Centre for Molecular Immunology, Havana, Cuba.

^2^ Metabolic and Cardiovascular Disease Laboratory, Group on Molecular and Cell Biology of Lipids, Alberta Diabetes and Mazankowski Heart Institutes, University of Alberta, Edmonton, AB, Canada.

^3^ Faculté de Pharmacie, Université de Montréal, Montréal, QC, Canada

^4^ Innovation and Managing Direction, Center for Molecular Immunology, Havana, Cuba

***Joint Corresponding Authors:**

Dr. Spencer D. Proctor, 4-002J Li Ka Shing Centre, University of Alberta, Edmonton, Alberta, Canada. T6G2P5.

Dr. Yosdel Soto. Centre for Molecular Immunology, 216 St and 15th Ave, Atabey, Playa PO Box 16040, Havana 11600, Havana, Cuba.

**SUPPLEMENTAL METHODS**

**Extracellular Matrix Preparation from Rat Smooth Muscle Cells**

Vascular smooth muscle cells (VSMC) were isolated using a tissue explant technique (51). Briefly, small pieces of the aorta were obtained from lean control rats and cultured in DMEM F12 containing 20% serum and Penicillin/Streptomycin. The cells isolated from lean rats were subsequently cultured in DMEM F12 containing 10% FBS and Penicillin/Streptomycin, NEAA, glutamine. In contrast, VSMC derived from IR rats were cultured in DMEM 25 mM of glucose in the presence of insulin (100 U). Cells between passages 5 to 10 were used for subsequent experiments. To produce VSMC-derived extracellular matrix (ECM), 10^5^ cells per well were cultured onto Nunc Lab Tek II Chambered slides until reaching confluence. The cells were lysed with 0.5% Triton X-100 in phosphate buffer (pH 7.4) for approximately 10 min followed by incubation with 25 mM NH_4_OH for 5 min to remove the cytoskeleton and the nuclei (51).

**Inhibitory Effect on Lipoprotein Binding**

Blocking experiments on lipoprotein binding to CS and VSMC-derived ECM *in vitro* were performed in similar conditions, as described for recognition experiments. In this case, the wells were pre-incubated with 125 μg/mL of chP3R99 mAb in sample buffer for 90 min at RT. Wells were incubated with Cy3-LDL or Cy5-RM (250 μg/mL). The specificity was assessed by using hR3 mAb as isotype-matched control. Finally, the the chamber separators were removed, and the slides were scanned at 550 nm and 620 nm (GenePix 4000B) to determine the fluorescence associated with the binding of LDL and RM, respectively.

**LDL Oxidation**

LDL oxidization was performed at a final concentration of 200 µg protein/mL in 20 µmol/L CuSO_4_ solution at 37°C for 18 h. The oxidation was verified by thiobarbituric acid reactive substance assay and ELISA using EO6 mAb. Native LDL or oxLDL conjugation to biotin was performed as described elsewhere (37).

**Biotinylated LDL Binding to Chondroitin Sulfate**

The binding of LDL to CS relative to oxidized LDL was assessed by ELISA. Immunoplates were coated with CS (10 µg/mL) in HEPES-buffered saline (HBS) and co-incubated with biotinylated LDL (0-2.5 µg/mL) in HBS containing 2 mmol/L CaCl_2_ and 2 mmol/L MgCl_2_.

The binding was detected with alkaline phosphatase-streptavidin complex (Jackson) and the absorbance measured at 405 nm. The samples were evaluated in triplicate for each condition and the coefficient of variation was <10%. Background values were less than 0.1.

**Immunogenicity of chP3R99 mAb in Rats**

The capacity of chP3R99 mAb (Ab1) to induce an idiotypic cascade of Abs in rats was evaluated by ELISA, as described in our previous studies (37). The kinetics of induction of anti-idiotypic Abs (Ab2 response) and anti-anti-idiotypic Abs (Ab3 response, autologous anti-CS rat Abs), both in IR and control rats immunized with this mAb. All the assays were performed in triplicate for each condition and the coefficient of variation was <10%. Background values were less than 0.1.

To detect the Ab2 response directed to chP3R99 mAb, Maxisorp (Nunc) ELISA plates were coated with chP3R99 or the control hR3 (10 μg/mL) overnight at 4°C in Carbonate/bicarbonate buffer, followed by a blocking step with 1% BSA in PBS for 1h at 37°C. The reactivity was detected by mean of a peroxidase-conjugated goat anti-rat serum (Jackson), incubated in blocking solution for 1h at 37°C. The reaction was revealed by using a TMB Liquid Substrate System for ELISA (Sigma-Aldrich) and the plates were read at 450 nm.

The kinetics of induction of autologous anti-CS Abs (Ab3 response), along with the reactivity of the sera to other GAGs and proteoglycans, was evaluated by ELISA as previously described (38). Briefly, Maxisorp (Nunc) plates were coated ON at 4ºC with GAGs or biglycan (10 µg/mL) in HEPES-buffer solution (HBS). After blocking with 1% BSA in HBS, the plates were incubated with the rat sera (1/400) for 90 min at RT in sample buffer. To specifically determine the IgG response, the plates were incubated with a peroxidase-conjugated goat anti-rat serum (Jackson), the reaction was revealed with a TMB substrate (Sigma-Aldrich), and the plates read at 450 nm.

All the assays were performed in triplicate for each condition and the coefficient of variation was <10%. Background values were less than 0.1.

**Tissue Homogenates**

Rat tissues (liver and aorta) were homogenized in lysis buffer [PBS (pH 7.4) with 1.5% Triton X-100 and 1% protease inhibitor cocktail (Sigma)] in liquid nitrogen. The samples were centrifuged at 17,000 × *g* for 10 min and the supernatants were stored at -80°C for further analysis. Total protein concentration was determined by BCA assay using a commercial kit and the concentration of the variables assessed was referred per mg of protein.

**Lipid Analysis**

High-performance liquid chromatography (HPLC) to quantify lipid concentrations in the liver and fast protein liquid chromatography (FPLC) fractions in plasma were also determined (Lipidomic core facility, University of Alberta, Canada). For lipoprotein fractioning, rat plasma (12 µL) was injected into an Agilent 1200 HPLC equipped with a Superose 6 gel filtration FPLC column. Cholesterol was visualized using in-line infusion of Infinity cholesterol reagent (Thermo Scientific, MA) and detection at 500 nm. For HPLC, lipids were extracted using a chloroform:methanol mixture (2:1), subsequently dried under nitrogen, and then reconstituted in chloroform:isooctane (1:1). A 10 µL aliquot of each extract was injected into an Agilent 1100 HPLC system equipped with an Onyx monolithic silica normal phase column. Detection was carried out using a Corona Ultra RS charged-aerosol detector. An internal standard of phosphatidyl dimethyl ethanolamine was used for each sample and the results were expressed in terms of lipid mass per milligram of protein.

**Insulin Resistance Assessments**

Glucose and insulin concentrations were quantified in plasma a week after the postprandial studies at the ending time point (ALPCO). The homeostasis model assessment of insulin resistance (HOMA-IR) score was calculated according to the formula: *Plasma insulin (µIU/L) * Blood glucose (mmol/L)/22.5*.

**Histological and Immunofluorescence Studies**

Histological studies were performed in fresh-frozen tissue sections with stained Oil red O-stained liver sections (10 µm). To assess the accumulation of chP3R99 mAb in different organs, tissue sections were incubated with a goat Cy3-conjugated anti-human IgG serum (Jackson) and counterstained with Hoechst. Images were digitally captured with a DP20 camera coupled to a fluorescence microscope Olympus BX51. And the images quantified blindly (ImageJ 1.50i).

**SUPPLEMENTAL RESULTS**

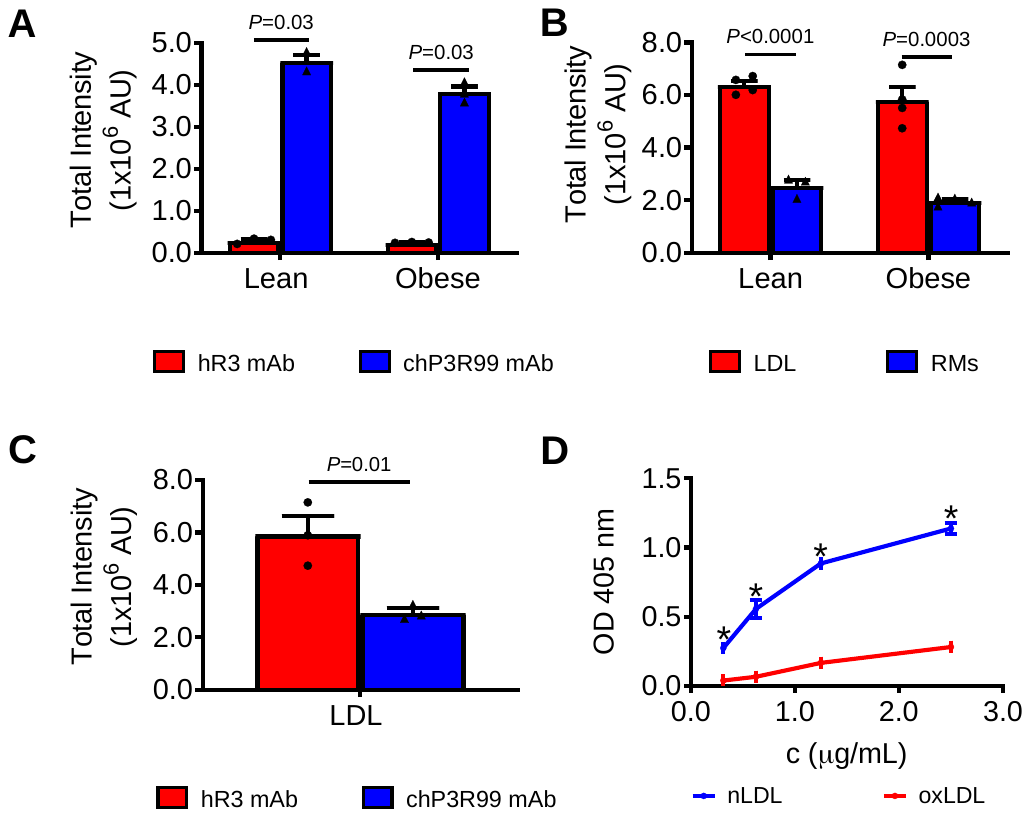

**Figure S1. Lipoprotein binding properties of VSMC-derived extracellular matrix and chondroitin sulfate *in vitro*.** **(A)** Recognition of chP3R99 mAb to ECM produced by VSMC from control rats cultured in normoglycemic conditions (5 mM) and ECM from VSMC from IR rats cultured in high-glucose media (20 mM) and insulin (100 U). ECM-coated Lab Tek II chamber slides were incubated with the mAbs at 50 μg/mL and the reactivity detected with a Cy3-conjugated goat anti-human IgG serum. **(B)** Lipoprotein binding to ECM produced by VSMC from lean controls or IR rats. Wells were incubated with Cy3-LDL or Cy5-RM at 250 μg/mL. **(C)** Inhibitory effect of chP3R99 mAb on LDL binding to vascular ECM produced by IR rats. Wells were pre-incubated with the mAbs (250 μg/mL) for 1h followed by incubation with Cy3-LDL diluted at 250 μg/mL. Slides were scanned at 520 nm and/or 640 nm to determine fluorescence associated with specific binding (GenePix 4000B). Specificity was assessed using hR3 mAb as isotype-matched control. **(D)** Binding of oxidized LDL to chondroitin sulfate relative to native LDL. ELISA plates were coated with 10 μg/mL chondroitin sulfate and incubated with biotinylated LDL or biotinylated oxidized LDL (0.31-2.5 µg/mL). The binding was detected with a streptavidine-alkaline phosphatase conjugate. Results are expressed as the mean±SEM of at least two experiments. Mann-Whitney *U* test or two-way ANOVA followed by Sidak multiple comparison test, **P*<0.05. RMs; remnants.

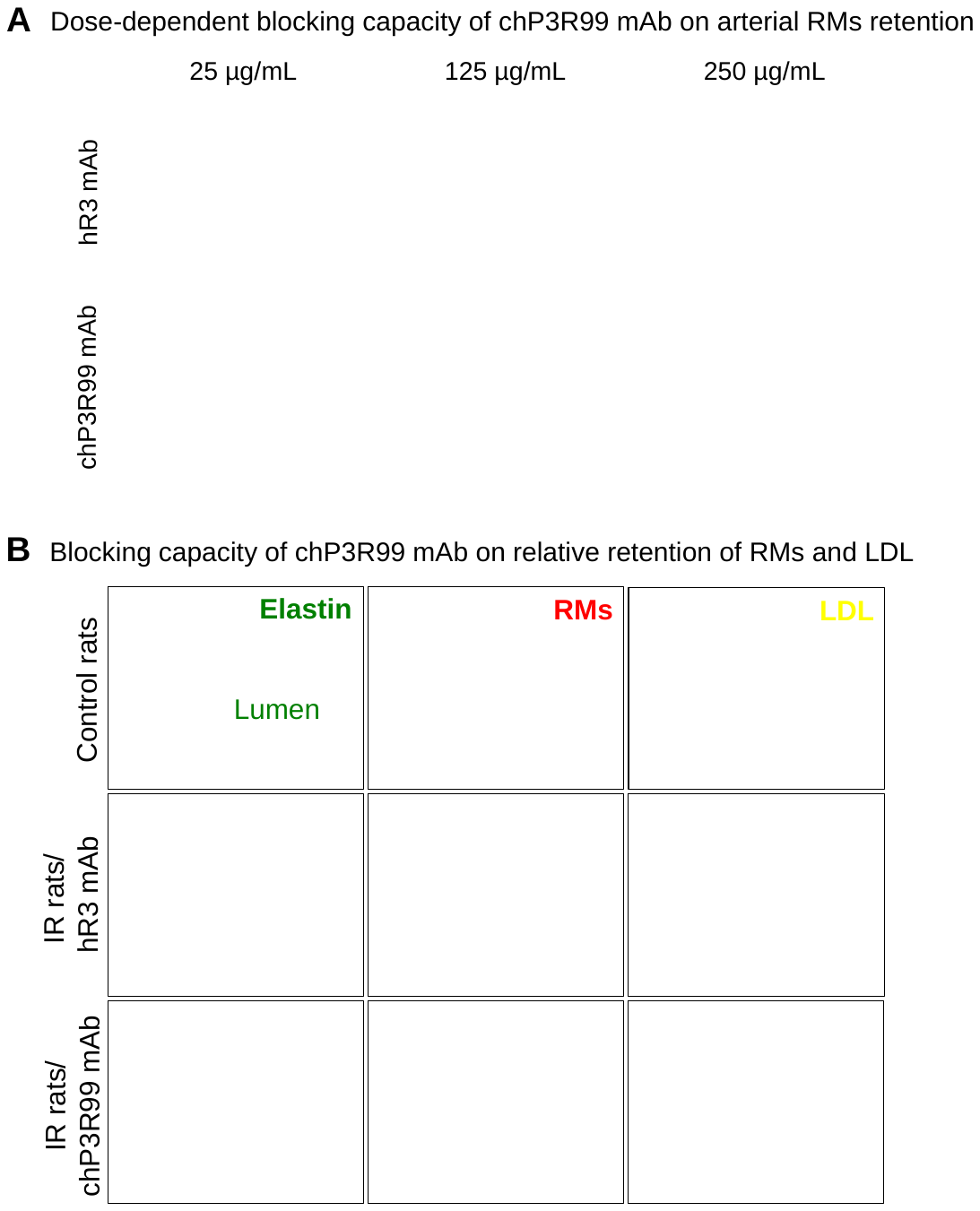

**Figure S2**. **Acute effect of chP3R99 mAb on arterial retention of remnants in insulin-resistant (JCR:LA-cp) rats.** Carotids from 12-weeks-old IR rats (n=4-6) were pre-perfused with chP3R99 mAb for 20 min followed by perfusion of fluorescently labeled lipoproteins. (A) Representative images showing a dose-response blocking effect of chP3R99 on the arterial retention of Cy5-RM (red) (150 µg/mL). (B) Representative images showing relative retention of lipoproteins in the carotids IR rats versus lean controls, along with the blocking effect of chP3R99 on Cy5-RM retention in the presence equivalent concentrations of Cy3-LDL (150 µg/mL). Bar =100 µm, Magnification 20X. Morphology is shown in green. Lean rats were perfused solely with lipoprotein preparation as control whereas specificity was assessed by using the hR3 mAb as isotype-matched control.

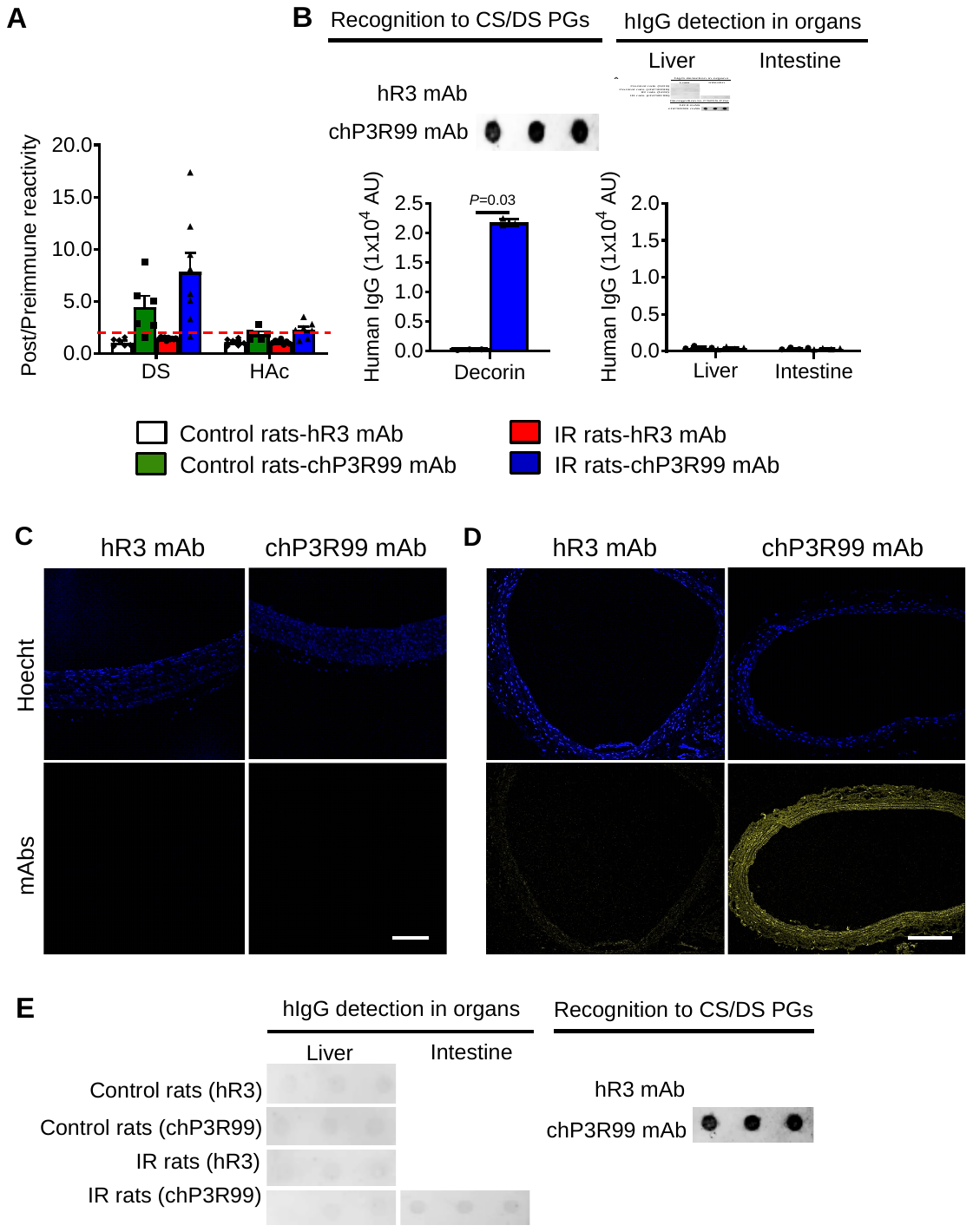

**Figure S3**. **Anti-GAG response in rats and tissue accumulation of the chP3R99 mAb.** **(A)** Ab3 anti-dermatan sulfate (DS) and anti-hyaluronic acid (HAc) response in rats immunized with chP3R99 mAb. ELISA plates coated with the GAGs (10 μg/mL) were incubated with the rat sera (1/400) followed by a peroxidase-conjugated goat anti-rat serum. Data are mean ± SEM, representing the ratio of the corresponding post-immune to pre-immune OD values. A cutoff of 2-fold was considered for the induction of Ab3 response (dotted line). **(B)** Tissue detection of chP3R99 mAb by Dot Blot in immunized IR rats. Membranes were spotted with decorin (20 µg/dot) and incubated with tissue homogenates (1 mg/mL). The mAbs were detected by a peroxidase-conjugated goat anti-human IgG secondary serum. Data are mean ± SEM. Mann-Whitney *U* test, **P*<0.05. **(C)** Representative images of chP3R99 detection (yellow) in IR rats immunized with chP3R99 mAb. carotid from IR rats perfused with chP3R99 were assessed as control **(D)**. The mAbs were detected by a Cy3-conjugated goat anti-human IgG secondary serum. Magnification 20X, Scale bar=100 µm. **(E)** Tissue detection of chP3R99 mAb by Dot Blot in rats treated with the mAbs in the acute setting. The experiment was conducted as described in **(B)**.

| **Table S1. Biochemical and Clinical Parameters in Control Rats Treated with chP3R99 mAb** | | | | | | | | | |
| --- | --- | --- | --- | --- | --- | --- | --- | --- | --- |
|  | **Acute injections (4.5 mg/kg, IV)** | | | | | **Immunizations (200 µg, SC, 6 wk)** | | | |
| **Parameter** | **hR3 mAb**  **(n=3)** | | **chP3R99 mAb**  **(n=3)** | | | **hR3 mAb**  **(n=5)** | | | **chP3R99 mAb**  **(n=6)** |
| Glucose, mmol/L | 7.0±0.6 | | 6.7±0.2 | | | 8.2±0.6 | | 9.1±0.5 | |
| Insulin, pmol/L | 120.2±12.7 | | 126.1±30.3 | | | 141.9±12.5 | | 136.9±10.7 | |
| HOMA-IR, AU | 11.6±5.6 | | 6.9±1.4 | | | 7.3±0.9 | | 7.7±0.8 | |
| Leptin, μg/mL | 5.2±0.6 | | 5.2±1.3 | | | 10.7±1.5 | | 13.1±3.5 | |
| Plasma LDL, mmol/L | 0.5±0.1 | | 0.6±0.1 | | | 1.0±0.3 | | 0.9±0.2 | |
| Plasma cholesterol, mmol/L | 1.7±0.2 | | 1.6±0.1 | | | 2.3±0.1 | | 2.1±0.1 | |
| Plasma triglycerides, mmol/L | 1.4±0.1 | | 1.1±0.1 | | | 0.3±0.1 | | 0.4±0.1 | |
| Liver cholesterol, µg/mgPr | 10.3±3.8 | | 8.5±1.1 | | | 6.0±0.8 | | 5.8±0.6 | |
| Liver triglycerides, µg/mgPr | 86.3±27.3 | | 80.7±17.7 | | | 44.8±6.3 | | 44.3±6.7 | |
| Aspartate aminotransferase, μg/mL | 17.8±0.2 | | 18.3±0.9 | | | 16.3±0.9 | | 14.9±1.3 | |
| Alanine aminotransferase, μg/mL | 4.1±0.6 | | 5.1±1.5 | | | 1.4±8.0 | | 2.0±0.6 | |
| Haptoglobin, mg/L | 2.7±0.4 | | 3.0±0.5 | | | 4.5±0.6 | | 3.9±0.8 | |
| C-reactive protein, mg/L | 236.1±18.1 | | 270.7±67.7 | | | 277.9±29.3 | | 268.8±17.2 | |
| β2-Microglobuline, μg/mL | 3.3±0.3 | | 3.6±0.4 | | | 3.7±0.5 | | 4.2±0.7 | |
| Results are mean±SEM. Two-tailed Mann-Whitney *U* test to compare chP3R99-receiving groups with their corresponding isotype-match controls. **P*<0.05, †*P*<0.01. HOMA-IR: Homeostatic Model Assessment for Insulin Resistance; mgPr: milligram of protein. | | | | | | | | | |
| **Table S2. Complete Blood Cell Counts in IR Rats Treated with chP3R99 mAb** | | | | | | | | | |
|  | | **Acute injections (4.5 mg/kg, IV)** | | | | **Immunizations (200 µg, SC, 6 wk)** | | | |
| **Parameter** | | **hR3 mAb**  **(n=3)** | | | **chP3R99 mAb**  **(n=3)** | **hR3 mAb**  **(n=5)** | **chP3R99 mAb**  **(n=6)** | | |
| WBC (x10^9^), cells/L | | 6.6±1.5 | | | 7.9±1.0 | 4.7±1.1 | 5.7±1.1 | | |
| RBC (x10^9^), cells/L | | 7.8±0.4 | | | 8.4±0.1 | 10.1±0.5 | 9.4±0.5 | | |
| Hemoglobin, g/L | | 133.8±2.7 | | | 136.3±1.8 | 169.7±7.3 | 157.9±8.8 | | |
| Hematocrit (x10^9^), cells/L | | 0.4±0.0 | | | 0.4±0.0 | 0.5±0.0 | 0.5±0.0 | | |
| Mean corpuscular volume, fL | | 55.2±3.1 | | | 52.6±0.4 | 49.3±1.0 | 48.8±0.7 | | |
| Mean corpuscular hemoglobin, pg | | 17.3±0.7 | | | 16.4±0.2 | 16.8±0.2 | 16.9±0.2 | | |
| Mean corpuscular hemoglobin concentration, g/L | | 342.3±5.3 | | | 345.9±5.6 | 342.3±5.3 | 345.9±5.6 | | |
| Cell hemoglobin concentration mean, g/L | | 315.8±2.7 | | | 316.3±1.5 | 329.3±3.4 | 323.6±6.1 | | |
| Mean cellular hemoglobin content, pg | | 17.3±0.8 | | | 16.5±0.1 | 16.2±0.1 | 15.7±0.1 | | |
| RBC distribution width, % | | 22.7±3.3 | | | 20.9±0.8 | 14.9±0.7 | 15.2±0.5 | | |
| Hemoglobin distribution width, g/L | | 28.1±1.2 | | | 25.6±1.0 | 26.1±0.9 | 26.3±0.5 | | |
| Platelets (x10^9^), cells/L | | 1150.0±227.6 | | | 1070.0±53.2 | 829.8±51.6 | 917.8±184.6 | | |
| Mean platelet volume, fL | | 7.7±0.2 | | | 7.6±0.0 | 11.1±2.2 | 15.5±2.7 | | |
| **WBC differential** | | | |  | |  |  | | |
| Neutrophils (x10^9^), cells/L | | 1.7±0.7 | | | 2.2±0.8 | 1.4±0.3 | 2.1±0.4 | | |
| Lymphocytes (x10^9^), cells/L | | 4.4±0.7 | | | 5.1±0.3 | 2.9±0.7 | 3.1±0.6 | | |
| Monocytes (x10^9^), cells/L | | 0.4±0.1 | | | 0.5±0.1 | 0.2±0.1 | 0.4±0.1 | | |
| Eosinophils (x10^8^), cells/L | | 0.4±0.2 | | | 0.7±0.4 | 0.7±0.1 | 0.9±0.5 | | |
| Basophils (x10^8^), cells/L | | 0.2±0.1 | | | 0.1±0.0 | 0.1±0.0 | 0.2±0.0 | | |
| Large unstained cells (x10^8^), cells/L | | 0.7±0.1 | | | 1.1±0.4 | 0.8±0.2 | 1.3±0.4 | | |
| WBC (Peroxidase method) (x10^9^), cells/L | | 5.8±1.0 | | | 8.8±1.0 | 5.6±1.3 | 7.2±1.4 | | |
| Reticulocytes (x10^9^), cells/L | | 713.9±182.5 | | | 566.5±50.4 | 204.0±4.8 | 223.0±24.2 | | |
| Lithium (x10^9^), cells/L | | 2.5±0.1 | | | 2.4±0.5 | 2.3±0.4 | 2.4±0.3 | | |
| Results are mean ± SEM. Two-tailed Mann-Whitney *U* test to compare chP3R99-receiving groups with their corresponding isotype-match controls. **P*<0.05, †*P*<0.01. WBC, White blood Cells; RBC, Red blood cells. | | | | | | | | | |
| **Table S3. Complete Blood Cell Counts in Control Rats Treated with chP3R99 mAb** | | | | | | | | | |
|  | | **Acute injections (4.5 mg/kg, IV)** | | | | **Immunizations (200 µg, SC, 6 wk)** | | | |
| **Parameter** | | **hR3 mAb**  **(n=3)** | | | **chP3R99 mAb**  **(n=3)** | **hR3 mAb**  **(n=5)** | **chP3R99 mAb**  **(n=6)** | | |
| WBC (x10^9^), cells/L | | 5.6±1.2 | | | 6.5±1.0 | 4.0±2.0 | 3.1±0.7 | | |
| RBC (x10^9^), cells/L | | 8.4±0.5 | | | 8.5±0.6 | 8.8±0.2 | 8.9±0.1 | | |
| Hemoglobin, g/L | | 146.6±4.8 | | | 143.0±6.4 | 156.0±4.4 | 162.8±2.3 | | |
| Hematocrit (x10^9^), cells/L | | 0.5±0.0 | | | 0.4±0.2 | 0.5±0.0 | 0.5±0.0 | | |
| Mean corpuscular volume, fL | | 53.4±1.6 | | | 51.9±1.3 | 52.8±1.6 | 52.4±2.4 | | |
| Mean corpuscular hemoglobin, pg | | 17.5±0.4 | | | 16.9±0.4 | 17.8±0.9 | 18.3±0.3 | | |
| Mean corpuscular hemoglobin concentration, g/L | | 337.3±8.9 | | | 350.5±11.6 | 337.3±8.9 | 350.5±11.6 | | |
| Cell hemoglobin concentration mean, g/L | | 333.2±2.9 | | | 332.8±2.8 | 327.7±4.2 | 333.0±6.2 | | |
| Mean cellular hemoglobin content, pg | | 17.7±0.4 | | | 17.2±0.3 | 17.3±0.3 | 17.4±0.6 | | |
| RBC distribution width, % | | 16.5±1.8 | | | 17.3±2.0 | 15.2±1.6 | 15.3±1.7 | | |
| Hemoglobin distribution width, g/L | | 23.8±0.4 | | | 25.0±0.3 | 21.6±1.3 | 22.6±1.0 | | |
| Platelets (x10^9^), cells/L | | 730.8±157.8 | | | 888.8±205.5 | 778.0±68.0 | 776.8±69.39 | | |
| Mean platelet volume, fL | | 9.5±0.8 | | | 8.2±0.8 | 12.0±4.2 | 14.2±3.8 | | |
| **WBC differential** | | | |  | |  |  | | |
| Neutrophils (x10^9^), cells/L | | 1.3±0.4 | | | 1.9±0.7 | 1.1±0.2 | 0.9±0.2 | | |
| Lymphocytes (x10^9^), cells/L | | 4.1±0.8 | | | 4.4±0.4 | 2.7±1.3 | 2.1±0.8 | | |
| Monocytes (x10^9^), cells/L | | 0.1±0.0 | | | 0.1±0.0 | 0.3±0.2 | 0.3±0.1 | | |
| Eosinophils (x10^8^), cells/L | | 0.7±0.2 | | | 0.6±0.2 | 0.5±0.1 | 0.4±0.1 | | |
| Basophils (x10^8^), cells/L | | 0.1±0.0 | | | 0.1±0.0 | 0.3±0.0 | 0.5±0.0 | | |
| Large unstained cells (x10^8^), cells/L | | 0.5±0.2 | | | 0.5±0.1 | 0.4±0.2 | 0.3±0.1 | | |
| WBC (Peroxidase method) (x10^9^), cells/L | | 6.6±1.6 | | | 7.4±1.3 | 4.4±1.9 | 3.7±0.7 | | |
| Reticulocytes (x10^9^), cells/L | | 406.3±105.8 | | | 447.4±130.3 | 152.2±18.4 | 121.6±7.9 | | |
| Lithium (x10^9^), cells/L | | 2.3±0.1 | | | 2.2±0.5 | 2.2±0.1 | 2.2±0.1 | | |
| Results are mean±SEM. Two-tailed Mann-Whitney *U* test to compare chP3R99-receiving groups with their corresponding isotype-match controls. **P*<0.05, †*P*<0.01. WBC, White blood Cells; RBC, Red blood cells. 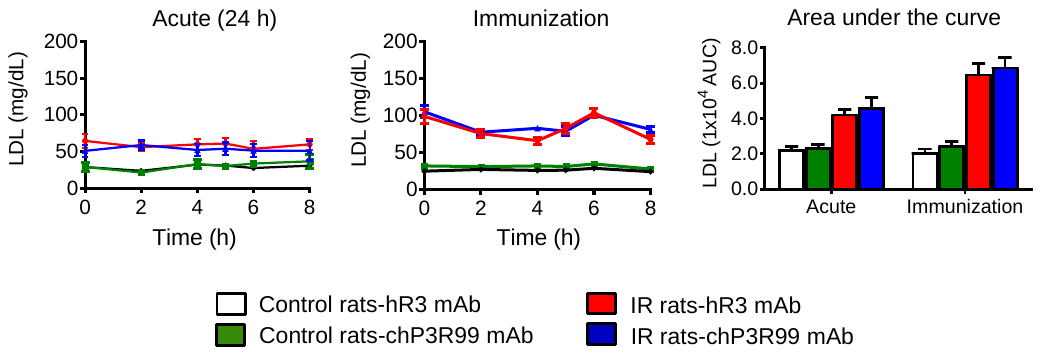 **Figure S4**. **Effect of chP3R99 mAb administration in postprandial LDL metabolism.** Lean control and cholesterol-fed IR rats received an acute intravenous administration of chP3R99 mAb (4.5 mg/kg) (n=3) or were immunized with six weekly subcutaneous injections of this mAb (200 µg) (n=6-7). LDL cholesterol was assessed 0-8 hours after a fat challenge and the area under the curve was determined in GraphPad Prism. Results are expressed as mean ± SEM. Student *t*-test, **P*<0.05. **Table S4. HPLC Lipid Quantification in the Liver of IR Rats Treated with chP3R99 mAb** \| \| \| \|; \| **Acute injections (4.5 mg/kg, IV)** \| \| **Immunizations (200 µg, SC, 6 wk)** \|; **Parameter*** **(µg of lipid/mgPr)** \| **hR3 mAb** **(n=3)** \| **chP3R99 mAb** **(n=3)** \| **hR3 mAb** **(n=4)** \| **chP3R99 mAb (n=4)**; Cholesterol esters \| 95.4± 13.8 \| 114.7±7.4 \| 78.0±11.6 \| 101.3±8.4; Free cholesterol \| 4.4±0.1 \| 4.2±0.3 \| 3.2±0.3 \| 4.2±0.2; Triglycerides \| 443.6±27.1 \| 605.8±27.6 \| 502.8±123.4 \| 759.9±80.0; Free fatty acids \| 17.9±7.5 \| 19.8±8.0 \| 13.5±3.8 \| 19.9±5.5; Phosphatidylcholine \| 39.9±0.9 \| 36.0±1.8 \| 37.9±5.0 \| 42.9±2.8; Phosphatidylethanolamine \| 18.4±1.4 \| 13.4±1.5 \| 14.7±2.7 \| 15.6±2.5; Phosphatidylinositol \| 4.5±0.8 \| 3.6±0.5 \| 3.9±0.9 \| 5.5±0.8; Phosphatidylserine \| 2.5±0.2 \| 2.2±0.3 \| 2.3±0.2 \| 2.8±0.2; Sphingomyelin \| 2.4±0.1 \| 2.4±0.2 \| 1.9±0.1 \| 1.7±0.1; Results are mean ± SEM. Two-tailed Mann-Whitney *U* test to compare chP3R99-receiving groups with their corresponding isotype-match controls. **P*<0.05, †*P*<0.01. mgPr, milligram of protein. (\| **Table S5. HPLC lipid quantification in the liver of control rats treated with chP3R99 mAb** \| \| \| \|; \| \| **Acute injections (4.5 mg/kg, IV)** \| \| **Immunizations (200 µg, SC, 6 wk)** \|; \| **Parameter** **(µg of lipid/mgPr)** \| **hR3 mAb** **(n=3)** \| **chP3R99 mAb** **(n=3)** \| **hR3 mAb** **(n=4)** \| **chP3R99 mAb (n=4)**; Cholesterol esters \| \| 6.3±3.4 \| 3.0±0.3 \| 4.4±2.6 \| 2.9±0.7; Free cholesterol \| \| 3.8±1.0 \| 3.0±0.2 \| 2.3±0.1 \| 2.7±0.3; Triglycerides \| \| 62.4±31.6 \| 36.3±5.6 \| 33.3±15.0 \| 25.45±4.9; Free fatty acids \| \| 22.6±0.2 \| 18.6± 8.7 \| 6.2±4.5 \| 13.1±2.9; Phosphatidylcholine \| \| 53.3±7.9 \| 63.3±6.8 \| 40.8±7.6 \| 60.3±7.3; Phosphatidylethanolamine \| \| 42.5±12.8 \| 29.4±2.4 \| 16.6±5.5 \| 29.2±3.5; Phosphatidylinositol \| \| 13.7±4.7 \| 8.6±1.3 \| 5.2±2.0 \| 8.3±1.3; Phosphatidylserine \| \| 3.4±0.7 \| 2.4±0.4 \| 2.3± 0.3 \| 3.1±0.3; \| Sphingomyelin \| 2.2±0.1 \| 2.5±0.2 \| 2.2±0.0 \| 2.1±0.1; \| Results are mean ± SEM. Two-tailed Mann-Whitney *U* test to compare chP3R99-receiving groups with their corresponding isotype-match controls. **P*<0.05, †*P*<0.01. mgPr, milligram of protein. \| \| \| \|) \| \| \| \| | | | | | | | | | |

| **Table S6. Cytokine and Growth Factors Levels in Aorta from IR Rats Treated with chP3R99 mAb** | | | | |
| --- | --- | --- | --- | --- |
|  | **Acute injections (4.5 mg/kg, IV)** | | **Immunizations (200 µg, SC, 6 wk)** | |
| **Parameter** | **hR3 mAb**  **(n=3)** | **chP3R99 mAb**  **(n=3)** | **hR3 mAb**  **(n=5)** | **chP3R99 mAb**  **(n=6)** |
| G-CSF (pg/mL) | ND | ND | ND | ND |
| GM-CSF, pg/mL | 4.4±0.1 | 4.7±1.0 | 3.9±0.5 | 5.1±0.5 |
| Eotaxin, pg/mL | 1.5±0.4 | 1.9±0.1 | 1.0±0.3 | 1.6±0.3 |
| MIP-1a, pg/mL | 54.9±1.3 | 52.3±1.0 | 50.9±2.3 | 51.9±3.2 |
| MIP-2, pg/mL | 132.8±32.4 | ND | 89.7±31.4 | 106.0±27.3 |
| MCP-1, µg/mL | 64.6±3.9 | 68.5±10.2 | 65.6±7.0 | 82.1±3.4 |
| Fractalkine, pg/mL | 13.3±1.3 | 24.0±7.1 | 12.2±2.5 | 15.4±5.4 |
| CCL5, pg/mL | 40.5±2.6 | 46.5±1.8 | 40.1±4.1 | 41.1±0.9 |
| LIX, µg/mL | 29.9±4.9 | 30.1±2.6 | 30.3±7.5 | 46.8±8.8 |
| IP-10, pg/mL | 7.5±4.3 | 5.5±2.5 | 4.1±1.2 | 4.2±1.9 |
| CXCL1, pg/mL | 15.1±0.9 | 17.5±3.9 | 34.6±11.4 | 27.1±1.9 |
| EGF, pg/mL | 26.1±13.7 | 13.2±6.7 | ND | 29.3±6.3 |
| VEGF, pg/mL | 151.6±22.0 | 113.3±26.4 | 181.7±38.1 | 140.8±3.2 |
| TGF-β1, µg/mL | 124.0±9.0 | 112.2±2.5 | 174.1±9.0 | 134.9±11.2 |
| TGF-β2, µg/mL | 127.9±14.2 | 108.6±9.2 | 158.3±29.5 | 144.5±28.5 |
| IFN-γ, pg/mL | 1117.0±39.6 | 1142.0±31.6 | 1110±17.0 | 1128.0±38.3 |
| TNF-α, pg/mL | 0.3±0.2 | 0.5±0.3 | 1.0±0.5 | 1.6±0.6 |
| IL-1α, µg/mL | 15.2±0.4 | 14.8±2.7 | 13.8±1.3 | 16.1±0.5 |
| IL-1β, pg/mL | 32.8±6.2 | 33.9±10.6 | 36.0±4.6 | 41.0±9.7 |
| IL-2, pg/mL | 17.2±1.4 | 15.1±6.3 | 15.6±2.9 | 19.9±1.3 |
| IL-5, pg/mL | 10.3±1.3 | 11.9±0.9 | 12.4±1.0 | 11.6±1.4 |
| IL-6, µg/mL | ND | ND | ND | ND |
| IL-10, pg/mL | 15.5±1.9 | 31.9±12.2 | 25.2±9.5 | 39.9±2.1 |
| IL-12p70, pg/mL | 25.6±2.8 | 35.3±4.9 | 20.5±3.8 | 22.5±3.3 |
| IL-13, pg/mL | 2.9±1.3 | 3.1±1.5 | 1.9±0.6 | 1.9±0.5 |
| IL-17A, pg/mL | 2.5±1.6 | 4.3±0.7 | 4.6±1.3 | 5.0±1.1 |
| IL-18, pg/mL | 306.6±23.6 | 358.4±37.9 | 367.7±28.4 | 387.5±14.3 |
| Results are mean±SEM. Two-tailed Mann-Whitney *U* test to compare chP3R99-receiving groups with their corresponding isotype-match controls. **P*<0.05, †*P*<0.01. ND, Not detected; G-CSF, Granulocyte colony-stimulating factor; GM-CSF, Granulocyte macrophage colony stimulating factor; MIP, Macrophage inflammatory protein; MCP, Monocyte chemoattractant protein; CCL5 (RANTES), C-C motif chemokine ligand 5; LIX, Lipopolysaccharide-inducible CXC chemokine; IP-10, Interferon gamma-induced protein 10; EGF, Epidermal growth factor; CXCL1 (GRO/KC), Chemokine (C-X-C motif) ligand 1; VEGF, Vascular endothelial growth factor; TGF, Transforming growth factor; IFN-γ, Interferon gamma; TNF-α, Tumor necrosis factor alpa; IL, Interleukin. | | | | |
